# Boundaries in the eyes: measure event segmentation during naturalistic video watching using eye tracking

**DOI:** 10.1101/2024.08.22.609279

**Authors:** Jiashen Li, Chen Zhengyue, Xin Hao, Wei Liu

**Affiliations:** Key Laboratory of Adolescent Cyberpsychology and Behavior (CCNU), Ministry of Education, Wuhan, China; Key Laboratory of Human Development and Mental Health of Hubei Province, School of Psychology, Central China Normal University, Wuhan, China

**Keywords:** event segmentation, naturalistic information processing, eye tracking, inter-subject correlation analysis, hidden Markov model

## Abstract

During naturalistic information processing, individuals spontaneously segment their continuous experiences into discrete events, a phenomenon known as event segmentation. Traditional methods for assessing this process, which include subjective reports and neuroimaging techniques, often disrupt real-time segmentation or are costly and time intensive. Our study investigated the potential of measuring event segmentation by recording and analyzing eye movements while participants view naturalistic videos. We collected eye movement data from healthy young adults as they watched commercial films (N=104), or online Science, Technology, Engineering, and Mathematics (STEM) educational courses (N=44). We analyzed changes in *pupil size* and *eye movement speed* near event boundaries and employed *inter-subject correlation analysis* (ISC) and *hidden Markov models* (HMM) to identify patterns indicative of event segmentation. We observed that both the speed of eye movements and pupil size dynamically responded to event boundaries, exhibiting heightened sensitivity to high-strength boundaries. Our analyses further revealed that event boundaries synchronized eye movements across participants. These boundaries, can be effectively identified by HMM, yielded higher within-event similarity values and aligned with human-annotated boundaries. Importantly, HMM-based event segmentation metrics responded to experimental manipulations and predicted learning outcomes. This study provided a comprehensive computational framework for measuring event segmentation using eye-tracking. With the widespread accessibility of low-cost eye-tracking devices, the ability to measure event segmentation from eye movement data promises to deepen our understanding of this process in diverse real-world settings.

## 1. Introduction

In fields such as cognitive psychology and neuroscience (including both human and animal subjects), experiments are commonly conducted on a trial-by-trial basis. However, our everyday life requires the processing of continuous information: we spontaneously segment continuous information into smaller units (i.e., event segmentation (Kurby & Zacks, 2008; Sun et al., 2020; Zacks, 2020; Zacks & Tversky, 2001)). Recently, there has been a growing interest among researchers in investigating cognitive processing during the continuous intake of information, such as watching movies (Hasson et al., 2004), sports (Antony et al., 2021) or listening to audiobooks (Yeshurun et al., 2017) and speech (Raccah et al., 2023). This focus is primarily due to the crucial role that event segmentation in continuous information processing plays in bridging perception and memory (Baldassano et al., 2017; Hasson et al., 2015; W. Liu et al., 2022). However, isolating and measuring event segmentation presents challenges given its simultaneous occurrence with various cognitive processes such as perception, attention, language understanding, and memory formation. In this study, we investigated whether human eye movements during continuous information processing, specifically during naturalistic video viewing, can serve as a novel signature for event segmentation.

Traditional methods of studying event segmentation, while informative, have their limitations. One common method to study event segmentation requires participants to mark event boundaries either during movie watching or retrospectively(Sasmita & Swallow, 2022). Despite its frequent use in prior studies (Kurby & Zacks, 2011; Zacks, Braver, et al., 2001; Zacks, Tversky, et al., 2001), this method has limitations: marking boundaries in real-time may alter the spontaneous ongoing segmentation process, and retrospective reporting may not accurately reflect immediate responses. Specifically, studies have shown that engaging in such overt segmentation tasks can even enhance subsequent memory for the observed events(Flores et al., 2017; Hanson & Hirst, 1989; Lassiter et al., 1988), possibly by increasing attentional resources devoted to important event changes. However, by requiring an explicit judgment, this method inherently alter the spontaneous nature of event segmentation as it unfolds during naturalistic viewing, shifting it towards a more goal-directed process. An alternate behavioral approach employs an independent cohort to determine event boundaries, which are then used to analyze another group’s data (Baldassano et al., 2017; Chen et al., 2017; Faber et al., 2018; Swallow et al., 2009). Nonetheless, individual performance on the segmentation task can vary significantly, potentially reflecting either true individual differences in segmentation or unaccounted systematic variances between groups(Sasmita & Swallow, 2022). Shifting focus to neural methods, researchers anticipate that neural activity may offer objective markers of event segmentation. Univariate analyses revealed that hippocampal activity increases near event boundaries during continuous experiences(Baldassano et al., 2017; Ben-Yakov et al., 2013; Ben-Yakov & Dudai, 2011; Ben-Yakov & Henson, 2018). Furthermore, multivariate analyses demonstrated that hippocampal and medial prefrontal regions form distinct event representations at these boundaries (W. Liu et al., 2022). These representational changes can be effectively captured using data-driven machine learning approaches (Baldassano et al., 2017). Nevertheless, the identification of neural-based event segmentation signatures still relies on predefined ’ground truth’ event boundaries, which remain contentious at the behavioral level for the reasons we discussed above. Moreover, neuroimaging is expensive and logistically complex to collect, particularly among special populations such as psychiatric patients, infants, and older adults. To further validate the correspondence between event segmentation at the self-report and neural levels, and to facilitate large-scale data collection across varied populations, it is crucial to identify an easily measurable signature of event segmentation that can be concurrently assessed alongside behavioral and neural responses.

In this study, we investigated an alternative method, notably eye tracking, to measure event segmentation during naturalistic video viewing, extending beyond conventional self-report (Sasmita & Swallow, 2022) and neural approach (Baldassano et al., 2017; Geerligs et al., 2021). Besides neural activity, previous studies already showed that continuous information processing can modulate various biological measures in humans, such as synchronizing heart rates across individuals (Pérez et al., 2021), but we selected the eye-tracking as our candidate measure for three reasons. First, eye tracking is capable of recording eye movements and pupil responses at the speed of cognition and neural activity (Kragel & Voss, 2022), and has been extensively used in research involving complex visual stimuli such as user interfaces (Lorigo et al., 2008), advertising (Bucher & Schumacher, 2006), and education (Madsen et al., 2021). Secondly, eye movement methods have been successfully combined with cognitive neuroscience techniques including fMRI, single-unit recording, and magnetoencephalography, allowing for more sophisticated investigations into memory that correlate discrete neural, memory, and eye movement events at fine temporal scale (Ryan & Shen, 2020). Lastly, in a sequence-learning task, Clewett and colleagues demonstrated that pupil dilation tracks changes in event structure (Clewett et al., 2020). Although this study did not use naturalistic videos but rather simple pictures and sounds to establish event structure, it highlights the potential of eye tracking to monitor complex event segmentation during naturalistic video viewing. Importantly, in Clewett’s research, due to the use of simple pictures as stimuli, only pupil responses were analyzed as signals of event segmentation. In our study, we concurrently consider pupil size and eye movement patterns including both their speed and trajectory. Previous research has demonstrated that eye movements show distinct patterns around event boundaries during video viewing. One study found that fixation durations decreased, while saccade amplitudes increased at event boundaries, and participants’ predictive gaze behaviors, such as fixations on objects about to be interacted with, declined around these boundaries (Eisenberg & Zacks, 2016). Moreover, two separate studies have highlighted increased gaze similarity across participants at event boundaries and noted that these gaze patterns significantly predict subsequent memory performance (E. E. Davis et al., 2021; Smith et al., 2024). Building on these foundational works, our study further investigates eye-tracking indicators of event segmentation.

In the analysis of pupil responses and eye movements elicited by naturalistic videos, our goal was to identify a signature of event segmentation using eye-tracking data. To this end, we went beyond the standard metrics of pupil diameter and movement speed, incorporating two advanced computational methods from neuroimaging into our eye-tracking analysis. The first method is the Hidden Markov Model (HMM) (Eddy, 2004), a recently introduced data-driven event segmentation model that was initially tested on fMRI data (Baldassano et al., 2017) and later confirmed using EEG data (Silva et al., 2019). The underlying assumption of the HMM is that discrete event representations can be treated as hidden states during the processing of a continuous narrative stimulus, where each event possesses a distinct signature that persists for the duration of the event. By fitting the HMM to eye-tracking data, which encompasses time series for pupil size and the x and y coordinates of fixation points, we can ascertain the optimal number of events as generated by the model, the characteristic eye-tracking pattern for each event, and the precise moments of event transitions. Subsequent studies have demonstrated that the HMM-based event segmentation model is sensitive to population characteristics, such as developmental changes(Cohen et al., 2022) and infant behaviors(Yates et al., 2022), as well as experimental manipulations, including anticipation(Lee et al., 2021) and schema(Masís-Obando et al., 2022). The second approach is the inter-subject correlation analysis (ISC)(Nastase et al., 2019), which quantifies the consistency of neural and biological responses among subjects. ISC was initially conceived to assess the correlation of fMRI responses to identical movie(Hasson et al., 2004) and robust correlations have since been observed across different data types, such as electroencephalogram (EEG)(Dmochowski et al., 2014), magnetoencephalography (MEG)(Lankinen et al., 2014), functional near-infrared spectroscopy (fNIRS)(Y. Liu et al., 2017), heart rate(Pérez et al., 2021), and eye movement(Madsen et al., 2021). Notably, studies have revealed that the degree of shared responses correlates positively with subsequent memory performance(Hasson, Furman, et al., 2008; Madsen et al., 2021; Meshulam et al., 2021). We proposed and tested the hypothesis that event boundaries can synchronize eye movements across participants. Consequently, ISC near these boundaries may serve as a distinctive signature for event segmentation. Moreover, a core function of event segmentation is to organize continuous experience into coherent, meaningful episodes. This principle can be directly tested using pattern similarity analysis (PSA), a method that quantifies the consistency of complex, multivariate patterns over time. In the context of the current study, PSA were applied to eye-tracking data. Specifically, we hypothesized that if the event boundaries are psychologically meaningful, then patterns of eye movements should exhibit higher similarity within the confines of an event than they do across event boundaries.

To overcome the inherent limitations of self-report and neural measures of event segmentation, this study systematically investigated the efficacy of eye-tracking-based metrics in quantifying event segmentation (**Figure 1**). We analyzed eye movement data from participants viewing two types of continuous naturalistic videos (i.e., commercial films, N=104 and STEM educational content, N=45). This design offers replication across two datasets and provides initial generalizability tests for different video types. First, we investigated the potential of both straightforward metrics (i.e., *pupil size* and *speed of moving*) and sophisticated computational techniques (i.e., *ISC* (Nastase et al., 2019) *and HMM* (Eddy, 2004)), to measure the event segmentation process. Specifically, we examined changes in these metrics before, during and after event boundaries and the correspondence of ISC and HMM-generated boundaries with human-annotated boundaries. Second, we explored how agreement on boundary locations among participants—termed boundary strength—affects these segmentation metrics. Third, recognizing the critical role of event segmentation in memory encoding, we investigated the influence of experimental manipulations on these indicators and their predictive power for subsequent memory performance. Ultimately, our goal was to identify an optimal eye-tracking signature for event segmentation that (1) is responsive to human-annotated boundaries, (2) sensitive to boundary strength and (3) experimental manipulation, (4) predictive of memory outcomes, and (5) generalizable across various naturalistic video stimuli.

**Figure 1.**
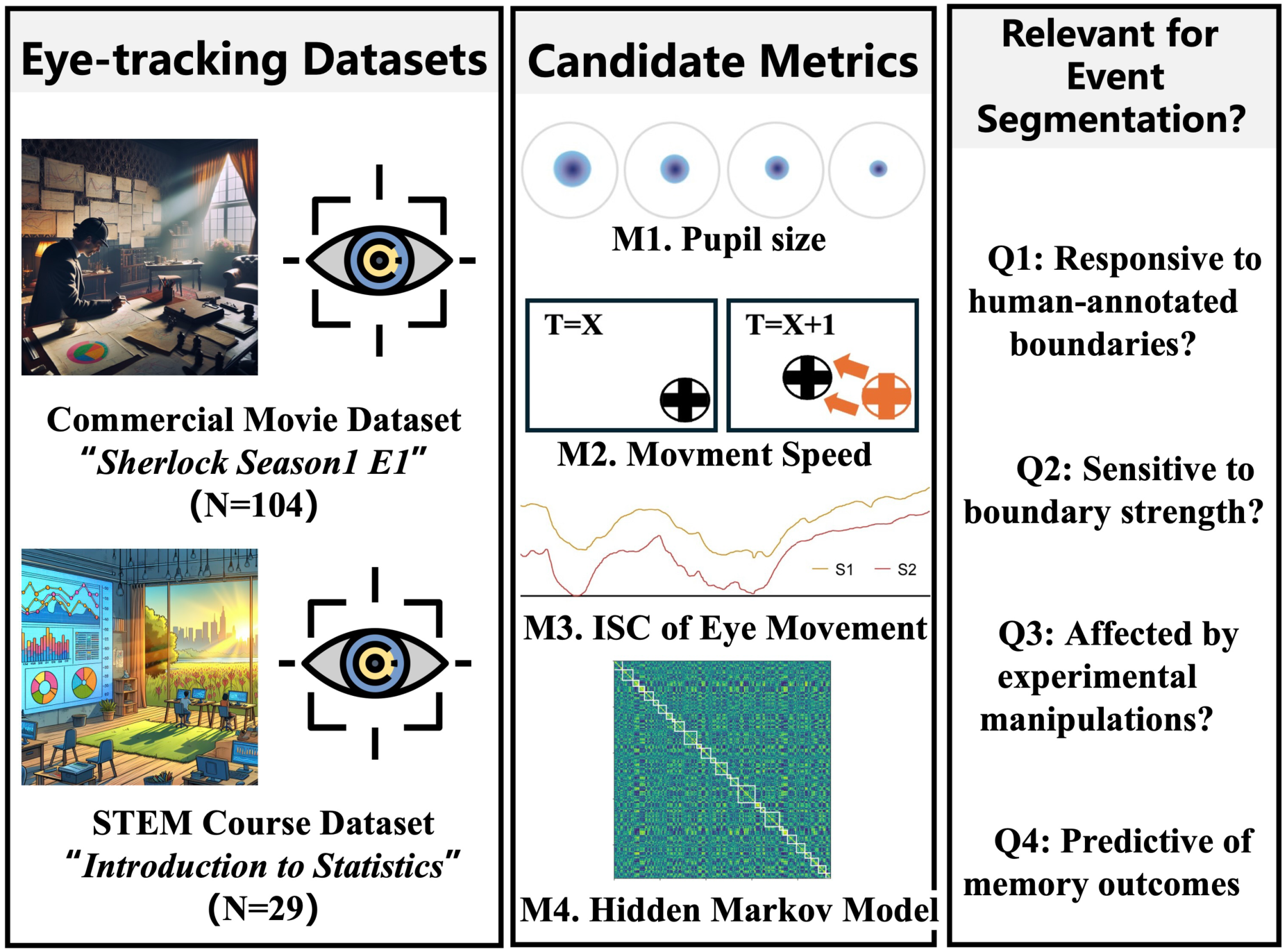
Analyzed Eye-Tracking Datasets, Candidate Metrics, and Analyses to Test the Relevance of Certain Metrics for Event Segmentation. We analyzed eye movement data from healthy participants (N=133) viewing either a commercial film, “*Sherlock Season 1, Episode 1*,” or an online educational clip, “*Introduction to Statistics* (Left Panel).” We examined the response of four eye-tracking-based metrics to event boundaries: pupil size, movement speed, Inter-Subject Correlation analysis (ISC), and Hidden Markov Model (HMM) (Middle Panel). For each metric in each dataset, we posed four questions to identify the optimal eye-tracking signature for event segmentation (Right Panel): (1) Is the metric responsive to human-annotated boundaries? (2) Is it sensitive to boundary strength? (3) Is it affected by experimental manipulation? (4) Is it predictive of memory outcomes?

## 2. Methods

### 2.1 Participants

#### Commercial Movie Dataset

A total of 112 right-handed, healthy university students (age: (M = 20.54) years, (SD = 2.04)) participated in this study. All participants had normal or corrected-to-normal vision, no hearing impairments, scores below 11 on the Beck Depression Inventory (BDI), indicating no current depression, and no history of neurological or psychiatric disorders. Furthermore, they confirmed having no prior exposure to the movie clip (i.e., Sherlock Season 1 Episode 1 “A Study in Pink”) used in this study. Recruitment was conducted via advertisements, and participants received compensation for their involvement. All participants provided written informed consent before the start of experimental procedure.

From the initial sample, eight subjects were excluded from the analysis due to six instances of data extraction failure and two programming errors, resulting in a final sample size of 104. Participants were randomly assigned to one of four groups to undergo different experimental manipulations (for details, see the experimental design section). Prior to viewing the film, the group assignments in the final sample were as follows: Short-R Group (N=30; 11 males, mean age = 20.32 years, SD = 2.08); Short-P Group (N=23; 8 males, mean age = 20.98 years, SD = 1.99); Long Group (N=25; 9 males, mean age = 20.18 years, SD = 2.11); and Schema Group (N=26; 12 males, mean age = 20.77 years, SD = 1.96). There were no significant differences among the groups in terms of age (F=0.57, p=0.63) or gender (χ²=0.88, p=0.82).

Notably, all participants viewed the same movie clip, and their eye movements were recorded using the same eye-tracking device. We aggregated the data to examine the general event segmentation process and conducted between-group analyses to understand how experimental manipulations affect event segmentation signatures. Experimental procedures were approved by the University Institutional Review Board for Human Subjects.

#### STEM Course Dataset

A total of 45 right-handed, healthy university students participated in this component of the study. All participants had normal or corrected-to-normal vision, no hearing impairments, and no history of neurological or psychiatric disorders. Participants watched a self-produced lecture tailored for STEM courses (i.e., Statistics) while their eye movements were tracked. Following the standard protocol, a subset of participants (N=11) was tasked with identifying and reporting event boundaries in the video. Additionally, an independent group of participants (N=15) viewed the video without undergoing memory tests or eye tracking. In total, data on eye-tracking and subsequent memory performance were collected from 29 participants, and 26 participants provided feedback on event boundaries. All participants affirmed they had not previously seen or learned the course material (i.e., Introduction to Statistics) in any format. Recruitment was executed through advertisements, with participants being compensated for their involvement. Experimental procedures were approved by the University Institutional Review Board for Human Subjects. All subjects provided written informed consent before the start of experimental procedure.

#### Sample Size Justification

As the first computational investigation to identify eye-tracking signatures for event segmentation during naturistic video viewing, precise effect size estimation prior to data collection was unfeasible. Our sample size was informed by prior studies on event boundaries, typically involving 20-30 participants and incorporating additional methodologies such as EEG and fMRI. A closely related study, which used pupil size to track event structures(Clewett et al., 2020), involved 34 participants and estimated a requirement of 25 participants to detect a large effect size (d=0.8) with an α of 0.05 and a power of 0.80. For each experimental group in our study, we aimed to collected at least 25 usable participants with complete memory and eye movement records. All our main and subgroup analyses involved more than 25 participants, except for the Short-P group, which included 23 participants following quality control procedures.

### 2.2 Materials and Design

#### 2.2.1 Commercial Movie Dataset

##### Experimental design

The experimental protocol was structured into three phases: the manipulation phase, the main video-watching phase, and the free recall phase. Initially, participants were instructed to watch approximately 45 minutes of video content attentively, with the objective of remembering the content for a subsequent memory test.

During the manipulation phase, the objective was to modulate the event segmentation process for the ensuing video-watching phase. In the Short-Random (R) group, participants watched 400 TikTok clips encompassing a variety of themes—humans, animals, objects, and scenes—with each category consisting of 100 clips. These clips, ranging from 1 to 5 seconds in duration, were pre-selected and sourced from a generic account to mitigate biases from the personalized recommendation algorithm. Conversely, participants in the Short-Personalized (P) group watched clips from their own accounts, suggested by personalized recommendation algorithms. The number of clips was not fixed, but the total viewing duration was maintained at 20 minutes, consistent with other groups. Additionally, the Long group viewed a 20-minute segment of the documentary “BBC Ocean,” and the Schema group watched a distinct 20-minute segment from the “Sherlock” film, which preceded the storyline of the main video content.

During the main viewing phase, all participants, regardless of group assignment, watched a 25-minute segment of “BBC Sherlock.” This choice was informed by its prevalent use in prior event segmentation research, known to elicit distinct event-specific neural representations, as evidenced by EEG and fMRI studies(Baldassano et al., 2017; Chen et al., 2017; W. Liu et al., 2022; Silva et al., 2019).

Following the main video-watching phase, during the free recall phase, participants were asked to provide a detailed verbal recount of the content viewed during the main phase, focusing on the richness of detail. Although the manipulation phase content was not directly assessed, participants were informed it would be tested to ensure sustained attention to the materials throughout the experiment. Video presentations during both phases were managed using Experiment Builder software (SR Research), and participants’ responses were initially recorded on smartphones before being transcribed for further memory analyses.

##### Human-labelled event boundaries and memory measures

We used the video stimuli from “Sherlock Season 1 Episode 1 ’A Study in Pink’”, which has been previously used in event segmentation studies(Baldassano et al., 2017; Chen et al., 2017; W. Liu et al., 2022), thereby we do not need to generate new event boundaries. The movie stimuli were presented in their original English language accompanied by both English and Chinese subtitles to facilitate comprehension for our native Chinese-speaking participants. Instead, we adopted the event boundaries delineated by Chen(Chen et al., 2017), encompassing 20 events within a 25-minute segment (i.e., 19 boundary locations). The audio recordings from the free recall phase were transcribed verbatim and subsequently assessed by two independent raters. Each participant’s recall was evaluated across three dimensions: (1) remember score: the quantity of events a participant could recall, (2) detail score: the comprehensiveness of details recalled for any given event, and (3) order: the accuracy of the event order recalled by the participant. Since order memory was not the primary focus of our study, remember and detail scores served as the principal metrics for memory performance, correlating with the eye-tracking-based event segmentation indexes. Initially, raters independently reviewed free recall responses. All free recall responses were scored by two independent raters who were blind to the experimental conditions. The raters used a pre-defined scoring sheet that specified the details and keywords for each event. Inter-rater reliability was calculated based on these initial, independent ratings, prior to any discussion. For the binary event recall (remembered/forgotten), reliability was excellent (Cohen’s Kappa = 0.91). For the number of details recalled, reliability was also high (Intraclass Correlation Coefficient, ICC(2,k) = 0.86). Following the initial rating, the two raters resolved any discrepancies through discussion to reach a 100% consensus. This final, consensus-based dataset was used for all subsequent analyses.

#### 2.2.2 STEM Course Dataset

##### Experimental design

The experimental protocol included both studying and testing phase. Before the beginning of the study phase, participants were briefed to watch the STEM learning material, specifically an introduction to statistics covering topics such as variables, constants, measuring scales, and the one-sample Z-test, under eye movement monitoring in anticipation of an immediate memory test thereafter. This STEM learning video, crafted by our lab members, lasts 12 minutes and 54 seconds. The STEM educational videos, presented in Chinese, did not include subtitles. The presentation of the video during the studying phase was managed by the Experiment Builder (SR Research). Following the video, the testing phase evaluated participants’ learning and memory across three dimensions (See details below). Notably, the ’remember score’ and ’test score’ from the memory tests served as the primary metrics for assessing video learning and correlated with eye-tracking data.

##### Human-labelled event boundaries and memory measures

To generate human-annotated event boundaries for this video, we recruited a sample of 26 participants who were tasked with identifying event boundaries and reporting them in minutes and seconds. Notably, boundary responses within a 3-second window were considered identical. It was generally observed that participants shared a consistent evaluation of event boundaries. Consequently, we selected 11 event boundaries corresponding to 12 distinct events for our main analyses based on the number of concepts taught in the video. This decision was made by the experimenters who prepared the course and slides. Additionally, we developed an index for each boundary, named “*boundary strength*,” to quantify the consensus on segmentation among participants. Event boundaries were categorized into three levels based on the degree of agreement: high strength (17 to 26 participants concurred), middle strength (2 to 16 participants concurred), and low strength (marked by only one participant). Each category included an equal number of eight boundaries to ensure comparable statistical power in subsequent analyses.

For the memory measure, all participant finished following memory tasks: (1) Image Recognition Task: Participants determined whether displayed images had appeared in the video amidst interference from similar internet-sourced images. This task’s performance was not analyzed in this study. (2) Free Recall Task: Participants wrote down key video points on a blank sheet. Performance was quantified as a ’remember score’. (3) Exam-like Memory Recall Task: Comprising retention and transfer tests, the retention test assessed direct recall of video content, while the transfer test measured comprehension through the application of learned concepts. Test items were derived from actual statistical exams to bolster ecological validity. The ’test score’ quantified performance.

### 2.3 Eye-tracking data acquisition

Participants in both the *Commercial Movie Dataset* and *STEM Course Dataset* were seated in the same room, with videos displayed on an identical monitor and eye movements tracked using the same device. To reduce the impact of lighting conditions on pupil size, the experimental room was shielded from outdoor sunlight with curtains, ensuring consistent luminance provided by indoor lighting regardless of the time of day. Eye movements were precisely monitored using a table-mounted, video-based EyeLink 1000 system (SR Research Ltd., Mississauga, Ontario, Canada) at a sampling rate of 1000 Hz, employing a 35-mm lens. Participants’ head movements were restricted throughout the experiment by a SR Research head and chin rest. Participants were informed about the eye movement calibration adjustment process and the setup of the eye tracker before starting experiments and were calibrated with the eye-tracker using a 9-point calibration grid.

### 2.4 Eye-tracking data preprocessing and analysis

#### Preprocessing

We used Matlab software to pre-process the eye movement data. Since the duration of the video material is known, we limited the length of the data sequence and made it uniform while ensuring its integrity. In addition, data points within these defined blink periods were subsequently replaced using linear interpolation for horizontal and vertical gaze coordinates and pupil size, and the complete data sequence of each subject was finally extracted, including screen fixation point X, Y and pupil size.

#### Pupil size and speed of moving

We extracted raw pupil size data from eye-tracking recordings and analyzed them without baseline correction, focusing primarily on dynamic changes within participants. The speed of movement was calculated using the x and y coordinates at the current and subsequent time points. Distances were computed using the Pythagorean theorem and subsequently converted to speed measurements. For each event boundary, we analyzed pupil size and movement speed within a 4-second window (i.e., from -2 to +2 seconds, with 0 representing the event boundary). To assess the effects of event boundaries on pupil size and movement speed, we conducted repeated-measures ANOVA to compare average pupil dilation responses and speeds at the time points of -2, -1, 0, 1, and 2.

#### Pattern similarity analysis of eye movements

To quantitatively assess whether the event boundaries identified by the human raters or our HMM-based approach resulted in segments with higher within-event similarity of eye-movement patterns compared to between-event similarity, we performed event-specific pattern similarity analyses. This provides a data-driven validation of the meaningfulness of the segmented events by examining the consistency of gaze behavior and pupil size within them. To analyze eye movement pattern similarity before and after event boundaries, we conducted a similarity analysis using eye-tracking data during video viewing. This analysis involved calculating similarity scores within 10-second intervals, initially based on pupil size and later including both pupil size and gaze coordinates (vertical and horizontal). Specifically, we derived a pupil size time series for each participant and computed correlations between two segments of data (50 points before vs. 50 points after the boundary) is calculated. The average of all these values provided an overall similarity measure for pupil size. This process was repeated for pupil size, vertical, and horizontal gaze. Finally, we computed a composite similarity measure by averaging the three similarity values, defined as similarity = (similarity_x_ + similarity_y_ + similarity_pupil_) / 3. Based on the selection of time window, we computed three types of similarity measures: (1) Between-event correlations were analyzed from -5 to 0 seconds and 0 to 5 seconds, with 0 indicating the event boundary. (2) Within-event post-boundary correlations were assessed from 0 to 10 seconds after the boundary. (3) Within-event pre-boundary correlations were analyzed from -10 to 0 seconds before the boundary. Statistical differences among these measures were evaluated using a repeated-measures ANOVA, which included event type as a three-level factor: pre-boundary, between-event, and post-boundary.

#### Interject-correlation analyses (ISC) of eye movements

To investigate the influence of event boundaries on the inter-subject correlation (ISC) of eye movements, we calculated the ISC for eye movements within specific temporal windows: before, during, and after an event boundary. Firstly, we calculated ISC within certain time window by 1) computing Pearson’s correlation coefficient between the vertical gaze positions of a single subject and those of all other participants as they viewed a video; 2) deriving an ISC value for each participant by averaging the correlation coefficients across all comparisons; and 3) replicating the above steps for every participant to obtain an individual ISC score per participant. We repeated this procedure for pupil size, vertical, and horizontal gaze. Subsequently, we computed a composite ISC score by averaging these three ISC values, defined as ISC = (ISC_x_ + ISC_y_ + ISC_pupil_) / 3. For each event boundary, ISC was evaluated under three different conditions: 10 seconds prior to the boundary (pre-boundary condition), 5 seconds before and after the boundary (boundary condition), and 10 seconds subsequent to the boundary (post-boundary condition). We then calculated the average ISC for each condition and participant, assessing differences using repeated-measures ANOVA.

#### Eye-tracking-based Hidden Markov Models (HMM)

We adapted the HMM approach previously used in fMRI(Baldassano et al., 2017) and EEG(Silva et al., 2019) to assess its efficacy in identifying latent eye-tracking states during video viewing, based on the event segmentation theory. Unlike fMRI and EEG data, the eye tracking method produced only three time series: X-axis and Y-axis coordinates of screen annotation points, and pupil size. Our objective is to deduce both the specific eye movement patterns and the structure of events, including the location of event boundaries. To this end, we reformulated our approach as a variant of the Hidden Markov Model. Specifically, we employed HMMs to identify event structures from continuous eye-tracking data using two distinct analytical approaches tailored to different research questions: (1) HMM for Data-Driven Event Boundary Discovery: For our primary analysis aimed at discovering event boundaries in a data-driven manner, the HMM was implemented with presetting a fixed number of events (k). In this approach, the model was trained on eye-tracking data from a randomly selected N-1 participants. The trained model, which identified an emergent number of states corresponding to distinct gaze patterns, was then applied to the hold-out participant’s data to predict event boundaries. These model-generated boundaries were subsequently compared to human-annotated event boundaries to assess model accuracy. (2) HMM for Assessing Individual Differences in Event Segmentation: To investigate individual differences in event segmentation granularity and its relationship with experimental manipulations and learning outcomes, we adopted a second HMM approach. For these analyses, we constrained the number of events (k) to a range of, for example 8 to 16. This range was selected to align with the conceptual level of event segmentation observed in human annotations, where an average of 12 distinct events were identified for the stimuli used. For each participant, the HMM was fit with varying numbers of states within this 8-16 range, and the optimal k was determined b subsequently. This approach allowed us to quantify individual variations in the number of segmented events around the human-perceived average and to test how these variations correlated with memory performance.

To evaluate the efficacy of the eye-tracking-based HMM in generating meaningful event structure, we employed two methods: (1) We assessed the fit of the eye-tracking data to the model by comparing both human-annotated and HMM-generated event boundaries using a within-event similarity measure with the null model. This measure’s group-averaged value was then tested against a null distribution, created by randomizing the event sequence 1000 times, to ascertain if it significantly exceeded the expected under random conditions. (2) We also compared the HMM-generated boundaries to human annotations by shuffling the event boundaries while preserving their duration to generate 1000 null distributions, and calculated a z-score that was transformed into a p-value to evaluate the significance of the match.

Furthermore, we investigated the performance of the HMM event segmentation across a range of k values (i.e., predefined number of events). Additionally, we examined whether individual differences in the model-defined optimal k values among participants signify significant variations in the event segmentation process. In the *Commercial Movie Dataset*, with an established ground truth of 20 events, we ran the HMM across k values from 15 to 25 and evaluated the resulting boundaries for their ability to model eye movement patterns, calculating within-event pattern similarity for each k and averaging these values across participants to assess whether the most accurate event structures yielded higher similarity scores. This analysis was also performed in the STEM dataset with the range of 8-16, because the ground truth is 12 events.

### 2.5 Pupil size correction for low-level confounds and gaze position

To ensure that our pupil-size analyses reflected cognitive processes rather than low-level sensory or motor artifacts, we implemented a correction procedure to remove confounding effects from the raw pupil-size signal. Following prior work, we identified four key potential confounds: screen luminance, motion, audio volume, and gaze position. For each participant’s data, we created time-series regressors for these confounds: Luminance: We calculated the mean grayscale luminance value across all pixels for each video frame and then downsampled it to match the sampling rate of the pupil data. Motion: We computed a frame-by-frame motion energy regressor by calculating the absolute difference in luminance values between consecutive frames. Audio Volume: We calculated the root-mean-square (RMS) of the audio signal amplitude in one-second windows, which served as a proxy for auditory volume. Gaze Position: To account for pupil foreshortening error, we included the horizontal (X) and vertical (Y) gaze coordinates at each time point as two separate regressors. These four time-series regressors (luminance, motion, volume, and X/Y gaze position) were then entered into a multiple linear regression model to predict the raw pupil size at each time point. The residuals from this regression model, which represent the pupil signal after accounting for these confounds, were retained as the “corrected pupil size”. All subsequent analyses reported in the main text were performed on this corrected pupil data to test the robustness of our findings.

### 2.6 Data and code availability

Processed behavioral and eye-tracking data, as well as custom-made video stimuli, are available at OSF (https://osf.io/n7k45/?view_only=673d89b23720410799b5daec1f5d511e)

Raw eye-tracking data is available upon reasonable request to the corresponding authors. Due to copyright restrictions, videos from the Commercial Movie Dataset are not accessible. Scripts for performing all eye-tracking analyses are located in the same repository as the data and stimuli. Data analyses were conducted using MATLAB or Python. Our HMM event segmentation model, was developed using the BrainIAK toolbox(Kumar et al., 2020) (https://github.com/intelpni/brainiak). Inter-subject correlation analyses were performed using in-house scripts, inspired by Pauline’ study(Pérez et al., 2021) (https://doi.org/10.17605/OSF.IO/MHVY7). Within-event and cross-event similarity analyses were conducted using in-house scripts, inspired by Silva’s study(Silva et al., 2019).

## 3. Results

### 3.1 Dynamics of pupil size and speed of eye movement in response to event boundaries

We investigated the changes in pupil size and the speed of eye movement around event boundaries, considering these dynamics as potential markers of event segmentation. Our analyses were first conducted with the *Commercial Movie Dataset* and was subsequently extended to the *STEM Course Dataset*. The latter serves as a conceptual replication, differing from a direct replication, due to the distinct characteristics of the two videos. Notably, the *STEM Course video* features pronounced visual transitions between slides and periods of static imagery while a scientific concept is explained.

When analyzing the *Commercial Movie Dataset,* we observed a pattern where pupil size increased immediately preceding the event boundary, peaking one second before (Max = 2108.2), followed by a sharp decrease shortly after the boundary (Min = 2080.54.7, F(4,412) = 9.137, *p*<0.001; **Figure 2a**). A corresponding pattern emerged for eye movement speed: an increase approaching the boundary, peaking at the boundary, and then a marked decrease thereafter (F(4,412) = 150.806, *p*<0.001; **Figure2b**).

When participants watched the *STEM course video*, pupil size decreased as it neared the event boundary, peaked at the boundary, and then reaching its lowest point a second after the boundary (Max = 978.516, Min = 937.125, F(4,112) = 13.392, *p*<0.001; **Figure2c**). This post-boundary reduction in pupil size aligns with the patterns noted in the *Commercial Movie Dataset*. The movement speed analysis for the *STEM Course dataset* echoed these findings, with the highest speeds recorded at the event boundary (F(4,112) = 41.576, *p*<0.001; **Figure2d**). Further analysis using a boundary strength measure was conducted exclusively on the *Commercial Movie Dataset* to explore how pupil size and movement speed varied with boundary strength. Specifically, we performed interaction analyses between category of boundary strength and time for pupil size (F(2,8)=1.256, *p*=0.266; **Figure 2E**) and movement speed (F(2,8)=1.971, *p*=0.049; **Figure 2F**). Pupil size responded significantly to high-strength boundaries (F(4,112)=4.866, *p*<0.001) and medium-strength boundaries (F(4,112)=2.957, *p*=0.023), but not to low-strength boundaries (F(4,112)=2.076, *p*=0.089). Movement speed responded significantly to high-strength boundaries (F(4,112)=3.657, *p*=0.008), but not to medium (F(4,112)=1.497, *p*=0.208) or low strength boundaries (F(4,112)=1.67, p=0.167). Our findings reveal that dynamic patterns in pupil size and movement speed in response to high-strength boundaries align with our main analyses. Conversely, responses to medium and low-strength boundaries are occasionally absent.

**Figure 2.**
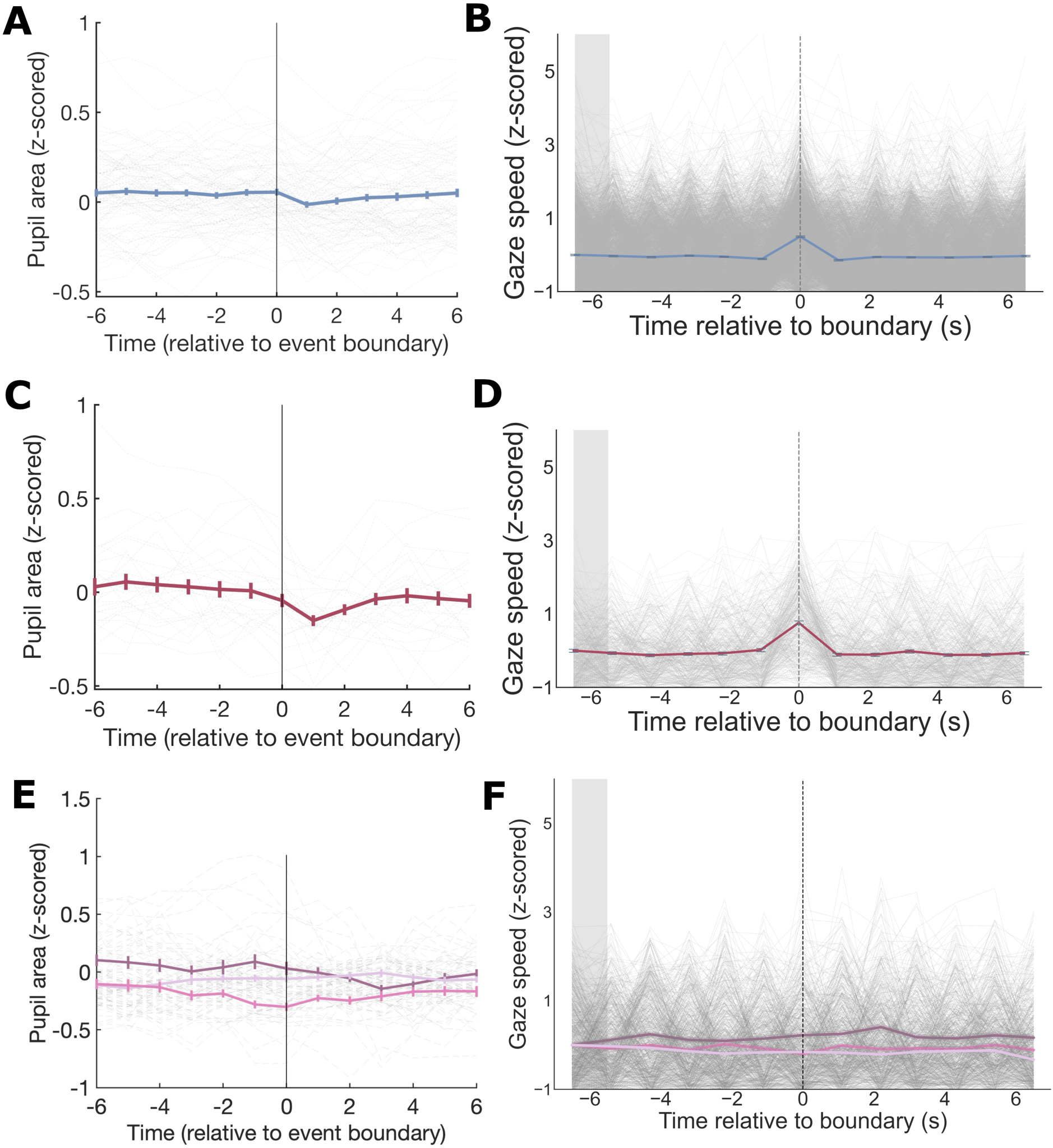
Pupil Size and Eye Movement Speed in Response to Event Boundaries and Their Modulation by Boundary Strength. (A) Pupil size dynamics around event boundaries in the Commercial Movie Dataset. (B) Eye movement speed dynamics around event boundaries in the Commercial Movie Dataset. (C) Pupil size changes around event boundaries in the STEM Course Dataset. (D) Eye movement speed dynamics around event boundaries in the STEM Course Dataset. (E) Impact of event boundary strength on pupil size adjustments in the Commercial Movie Dataset. (F) Influence of event boundary strength on eye movement speed patterns in the Commercial Movie Dataset.

### 3.2 Eye movement pattern similarity within and across event boundaries

Event Segmentation Theory suggests that mental representations within an event are more stable than those across events, indicating dynamic changes in representations at detected boundaries. This concept is supported by EEG neural pattern similarity analyses(Silva et al., 2019). We explored this prediction using eye-tracking data, initially with the *Commercial Movie Dataset* and subsequently with the *STEM Course Dataset*. Specifically, we conducted a point-to-point similarity analysis of eye-tracking segments from −10 to +10 seconds around the boundary time point, using pupil size or together with vertical and horizontal gaze coordinates. We performed this analysis between three distinct pairs of temporal intervals: the within-event pre-boundary condition (−10 to −5 s and −5 to 0 s relative to the boundary), the between-event condition (−5 to 0 and 0 to 5 s relative to the boundary), and the within-event post-boundary condition (0 to 5 s and 5 to 10 s relative to the boundary). We then calculated the average similarity values for each condition and subject, with differences assessed using repeated-measures ANOVA.

In the *Commercial Movie Dataset*, analyses of pupil size alone revealed no significant differences in similarity measures across within-event pre-boundary, between-event, and within-event post-boundary conditions (F(2,206) = 2.264, *p* = 0.107; **Figure 3A**). However, when pupil size was analyzed in conjunction with vertical and horizontal gaze, a higher similarity measure was observed for the between-event condition (F(2,206) = 3.676, *p* = 0.027; **Figure 3B**), indicating that combining eye movements with pupil size more effectively captures event segmentation processes.

**Figure 3.**
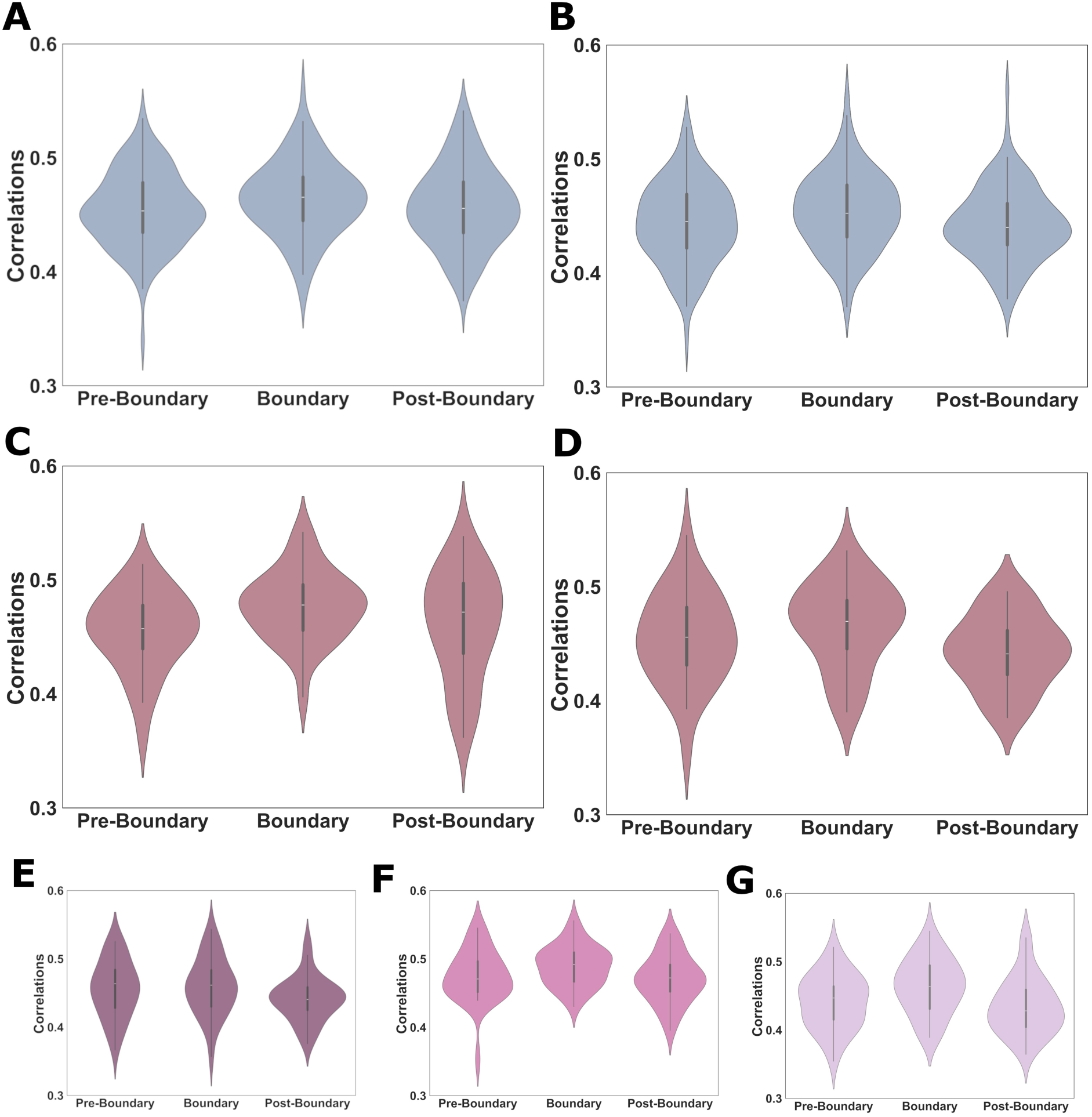
Eye movement pattern similarity within and across event boundaries during continuous video watching. (A) We analyzed three similarity measures derived from pupil size—within events before the boundary, across events separated by boundaries, and within events after the boundary—using the *Commercial Movie Dataset*. (B) Higher similarity measure, incorporating both pupil size and vertical and horizontal gaze, was calculated in the across-event condition, compared to two within-event conditions, using the *Commercial Movie Dataset*. (C) Participant’s three similarity measures derived from pupil size for three event conditions, analyzed with the *STEM Course Dataset*. (D) Participants’ similarity measures, calculated from both pupil size and vertical and horizontal gaze, were evaluated for three event conditions with the *STEM Course Dataset*. (E–G) Combined similarity measures (pupil size and gaze position) stratified by event-boundary strength—high (E), medium (F), and low (G)—in the STEM Course Dataset.

Replication analyses using the *STEM Course Dataset* produced similar, non-significant results for analyses of pupil size alone (F(2,56) = 2.534, *p* = 0.088; **Figure 3C**) and in combination with gaze (F(2,56) = 2.861, *p* = 0.066; **Figure 3D**). Notably, no interaction was found between boundary strength and time in the analyses of pupil size alone (F(2,4) = 0.529, *p* = 0.715). However, an interaction was observed in the analysis combining pupil size with gaze metrics (F(2,4) = 3.385, *p* = 0.011). Specifically, when analyzing high-strength event boundaries, there was a significant increase in similarity measures between low-strength boundaries (F(2,56) = 3.822, *p* = 0.028; **Figure 3G**). However, this was not observed for high-strength (F(2,56) = 1.756, *p* = 0.182; **Figure 3E**) or middle-strength boundaries (F(2,56) = 2.515, *p* = 0.09; **Figure 3F**).

### 3.3 Eye movement synchrony across participants at event boundaries

We applied inter-subject correlation (ISC) analysis to two eye-tracking datasets to assess whether and how eye movements synchronize across participants in response to an event boundary during continuous video viewing. Using both vertical and horizontal gaze coordinates along with pupil sizes, we analyzed ISC across three conditions (**Figure 4A**): 10 seconds before the boundary (pre-boundary condition), 5 seconds before and after the boundary (boundary condition), and 10 seconds following the boundary (post-boundary condition).

**Figure 4.**
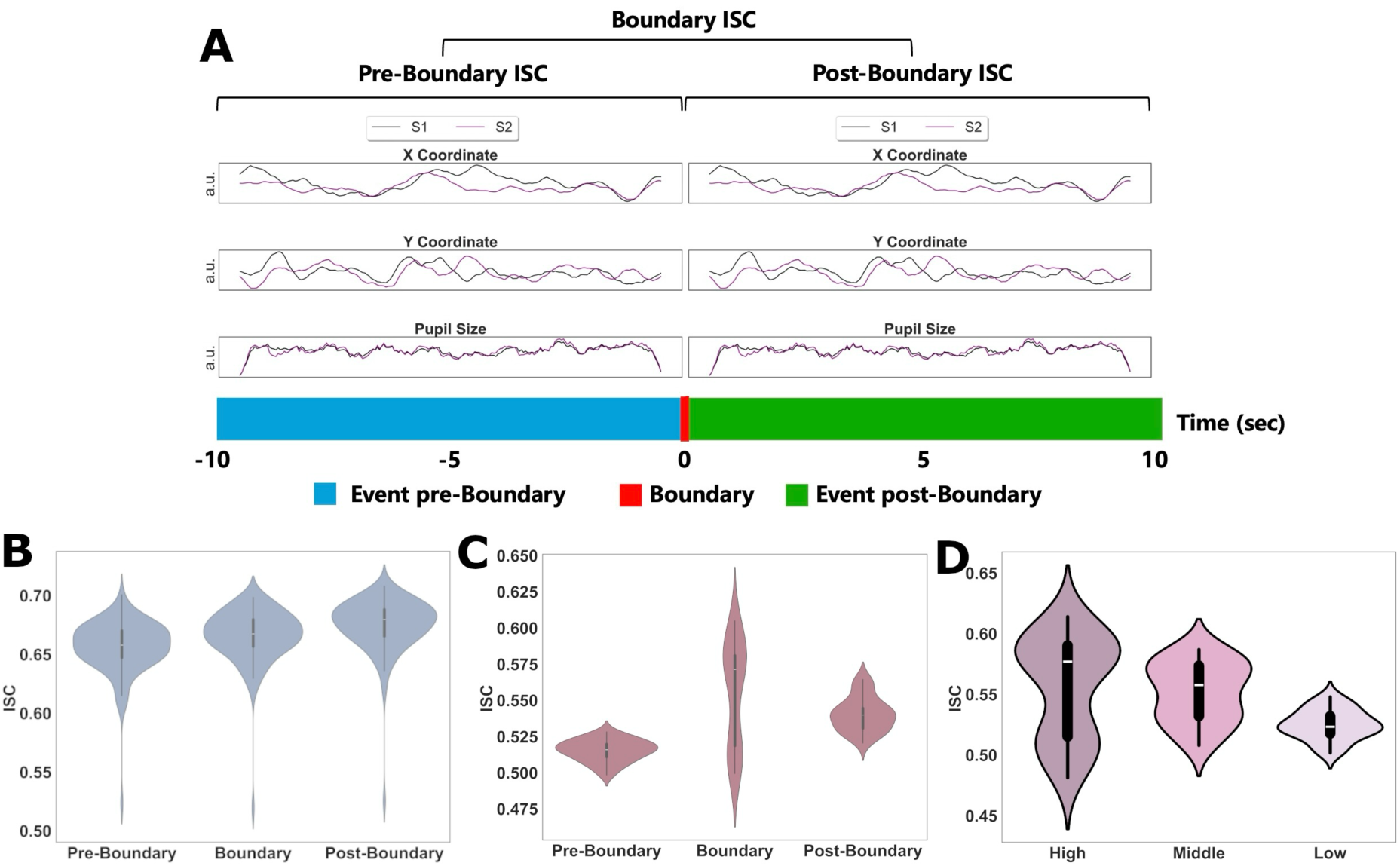
Synchronized Eye Movements in Response to Event Boundaries. **(A)** Synchronized eye movements—specifically, the positions (X and Y coordinates) and pupil sizes—of two participants (S1 and S2) while continuously watching two distinct events separated by an event boundary. Intersubject correlation (ISC) is quantified as the mean correlation between both vertical and horizontal gaze positions and pupil sizes across participants. We analyze three ISC metrics: pre-boundary ISC, calculated during the final 10 seconds before the event boundary; boundary ISC, derived from 5 seconds of eye-tracking data from each event; and post-boundary ISC, measured during the initial 10 seconds following the event boundary. **(B)** Comparison of three kinds of ISC within the *Commercial Movie Dataset*. **(C)** Comparison of three kinds of ISC within the *STEM Course Dataset*. **(D)** Comparison of boundary ISC associated with event boundaries characterized by high, medium, and low boundary strengths.

In the *Commercial Movie Dataset*, distinct ISC patterns emerged for three different ISCs (F=81.16, *p*<0.001; **Figure 4B**): boundary ISC is significantly higher than pre-boundary ISC (t=5.37, *p*<0.001, Cohen’s d=0.52), with a continued increase to post-boundary ISC (t=6.88, *p*<0.001, Cohen’s d=0.67). Similarly, in the *STEM Course Dataset*, we confirmed significant ISC variations across conditions (F=29.28, *p*<0.001; **Figure 4C**): boundary ISC is significantly higher than pre-boundary ISC (t=5.96, *p*<0.001, Cohen’s d=1.11), but it decreased post-boundary (t=-2.55, *p*=0.017, Cohen’s d=-0.47), though remaining higher than baseline level (i.e., pre-boundary) (t=11.31, *p*<0.001, Cohen’s d=2.1).

Additionally, we demonstrated that boundary strength influenced boundary ISC levels (F=22.48, *p*<0.001; **Figure 4D**): high and middle strength boundaries led to greater synchronization of eye movements than low strength boundaries, as shown by significant t-tests (t=4.81, *p*<0.001, Cohen’s d=0.89 for high strength; t=7.33, *p*<0.001, Cohen’s d=1.36 for medium strength), suggesting that stronger boundaries are more likely to induce synchronized eye movements across participants during event boundaries.

### 3.4 Discovering event structures in continuous eye-tracking recordings from humans

We employed two approaches to test the hypothesis that meaningful event structure information can be extracted from eye-tracking data using a Hidden Markov Model (HMM) event segmentation model when participants watched continuous video. First, we examined whether and to what extent the eye-tracking data fitted the model based on human-annotated or HMM-generated event boundaries. To do this, we conducted the similarity analysis of the eye-tracking data at each time point during video watching and calculated the degree of similarity within each event. We then tested this group-averaged similarity value against a null distribution generated by running the same analysis 1000 times with a shuffled temporal order of events. Second, we employed the HMM-based event segmentation method to automatically discover event-tracking-based signatures and temporal boundaries in the dataset. Then, we validated our model by comparing HMM-generated event boundaries to human annotations.

In the *Commercial Movie Dataset*, we firstly showed a higher degree of similarity values within events from the human-annotated event boundary model compared to correlation value from their null distribution at the group level (*p*<0.001; **Figure 5B**). This is also true for event boundaries generated by HMM (*p*<0.001 ; **Figure 5C**). Next, we then measured, what fraction of its HMM-generated boundaries were close to human annotations. At the group level, the HMM-generated boundaries showed an above-chance match to human annotations (*p*<0.001; **Figure 5D**).

**Figure 5.**
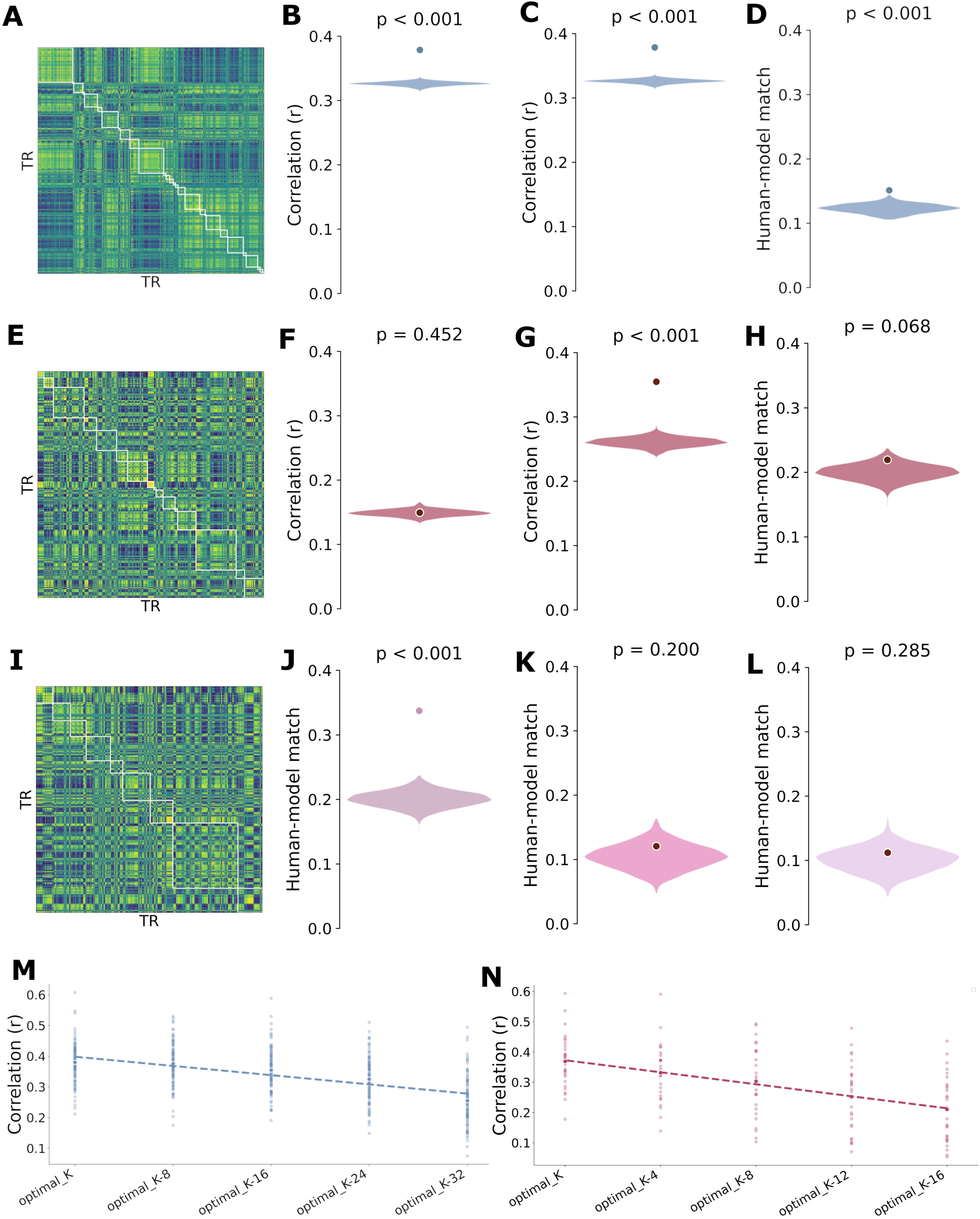
Hidden Markov Model (HMM)-based Event Segmentation Model for Eye-Tracking Data. Analyses on the Commercial Movie dataset (A-D, M): (A) Example alignment of HMM and human-annotated event boundaries. (B-C) Within-event gaze similarity was significantly higher than the null distribution (histograms) for both (B) human-annotated and (C) HMM-generated boundaries. The light blue circle indicates the observed participant average. (D) HMM-human boundary matching was significantly above chance. Corresponding analyses on the STEM Course dataset (E-L, N): (E) Example boundary alignment. (F-G) Within-event similarity for (F) human and (G) HMM boundaries. (H) HMM-human boundary matching. Boundary strength and model parameter analyses: (I) Example alignment for a high-strength boundary in the STEM dataset. (J-L) Boundary matching was modulated by strength, with the HMM effectively identifying high-strength but not medium-or low-strength boundaries. (M-N) The relationship between the HMM’s optimal event count (K) and gaze similarity is shown for the (M) Commercial Movie and (N) STEM Course datasets.

In the *STEM Course dataset,* however, we did not find significant higher degree of similarity values within events from the human-annotated event boundary model compared to correlation value from their null distribution at the group level (*p*=0.452; **Figure 5F**). Instead, we found the eye-tracking data fit the model generated by HMM (*p*<0.001 ; **Figure 5G**), suggested by a higher degree of similarity compared to null model. Secondly, the HMM-generated boundaries showed marginally above-chance match to human annotations (*p*=0.0688 ; **Figure 5H**). However, when we examined the influence of boundary strength on the congruence of HMM-defined and human-annotated boundaries, we found that the HMM was more effective in identifying boundaries of high strength compared to those of middle and low strength. The HMM pinpointed the optimal timing for events with high-strength boundaries (*p*<0.001; **Figure 5J**), but not for events with medium (*p*=0.200; **Figure 5K**) or low-strength boundaries (*p*=0.285; **Figure 5L**).

The HMM event segmentation model can autonomously determine the optimal number of events within time series data. We aimed to use the model-derived optimal event number (i.e., the k value) to represent individual differences in the event segmentation process. Before assessing the cognitive significance of the k value, we validated it using similarity analyses. Within a specified range of candidate k values (See Methods for details), we instructed the HMM to rank these k values from most to least optimal and used various k values along with its corresponding boundary locations to compute similarity metrics within our eye-tracking data. We observed that the application of the optimal k value yielded the highest similarity values. Moreover, as the k value deviated from the optimal, the similarity scores decreased significantly. This trend was consistent across both the *Commercial Movie Dataset* (*p*<0.001; **Figure 5J**) and the *STEM Course dataset* (*p*<0.001 ; **Figure 5K**).

### 3.5 HMM-based event segmentation are sensitive to experimental manipulations and predict learning outcomes

To assess the sensitivity of our HMM-based eye-tracking measures to variations in stimulus characteristics and potential viewing-history effects, we compared event segmentation patterns across participants exposed to different types of video content. This comparison allows us to demonstrate the utility of our method in detecting subtle but systematic differences in how viewers segment ongoing experiences based on stimulus properties and prior viewing habits, which is a critical aspect for a methodological paper aiming to offer a robust tool for future event segmentation research. Our study used the *Commercial Movie Dataset* to investigate this validation. Participants were divided into two experimental subgroups who viewed short TikTok video clips (Short-R for random videos and Short-P for personalized videos) before watching a continuous movie clip. Two control groups included participants who watched a movie-unrelated documentary clip (Long group) or a movie-related movie clip (Schema group).

We hypothesized that exposure to short video clips might impair subsequent event segmentation during movie watching, as indicated by altered eye-tracking-derived event segmentation metrics. The analysis focused on assessing differences in the individual optimal K value and p value, reflecting whether eye-tracking data fit the model based on HMM-generated event boundaries. After determining the optimal K value for each participant, we compared K values across groups and found a significant group difference (F=7.25, *p*<0.001; **Figure6A**). Notably, watching randomly selected TikTok videos, compared to the Long group, resulted in higher K values (t=2.18, *p*=0.034, Cohen’s d=0.28), suggesting the more fragmented event segmentation during video watching. Contrary to our prediction, watching personalized TikTok video clips (Short-P) before movie watching, compared to the Long group, resulted in significantly lower K values (t=-2.9, *p*=0.006, Cohen’s d=-0.31). Additionally, we analyzed p-values as an index for event segmentation and found the effect of experimental manipulations before movie watching on p-values (F=4.14, *p*=0.008; **Figure 6B**). The two experimental groups did not differ from each other (t=0.71, *p*=0.48, Cohen’s d=0.19). Both the Schema and Long group demonstrated lower p values compared to the Short-P (Schema: t=-2.69, *p*=0.01, Cohen’s d=-0.77; Long: t=-2.32, *p*=0.02, Cohen’s d=-0.67) and Short-R (Schema: t=-2.62, *p*=0.01, Cohen’s d=-0.70; Long: t=-2.13, *p*=0.03, Cohen’s d=-057) group.

**Figure 6.**
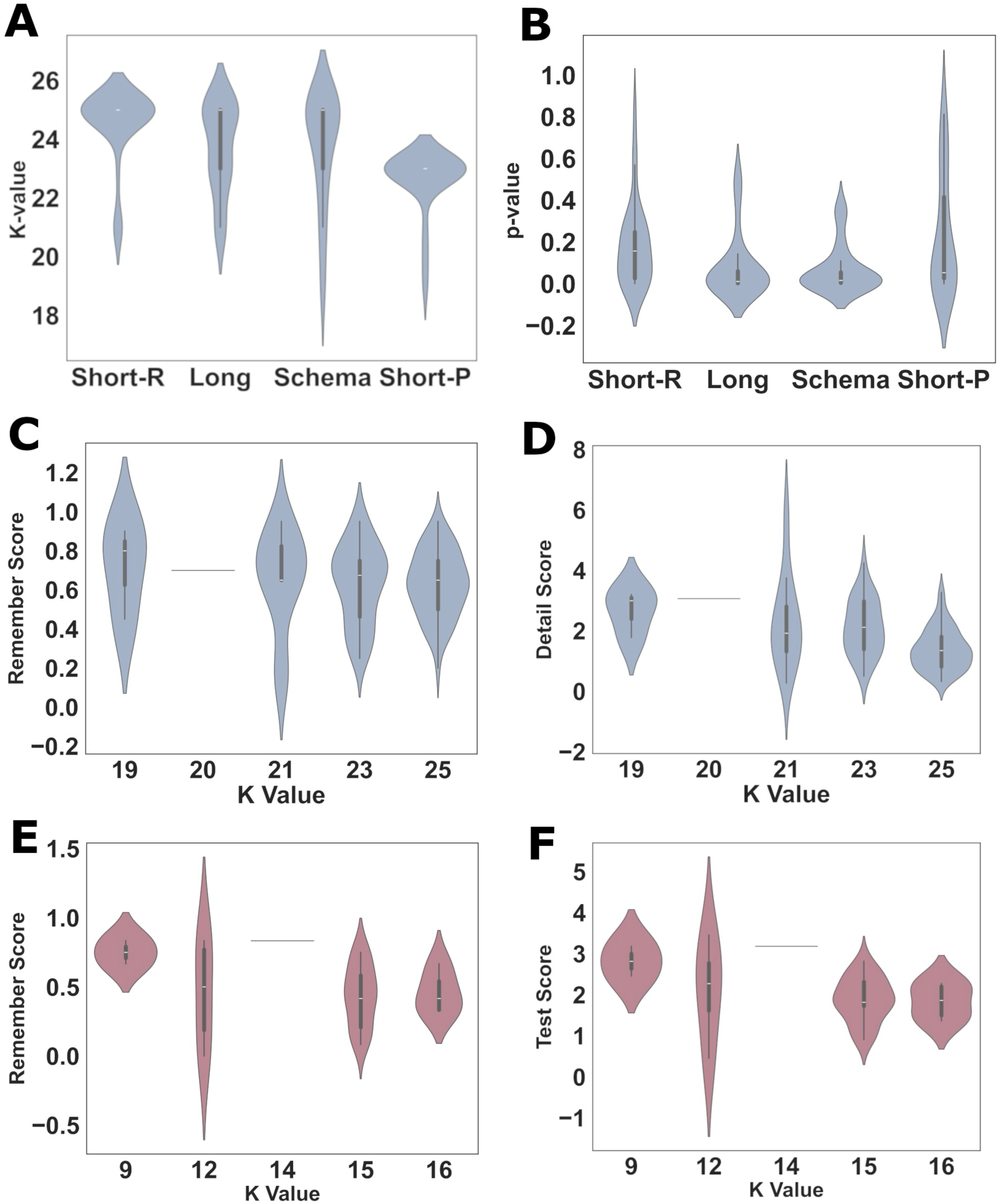
Validity of HMM-Based Event Segmentation Measures: Sensitivity to Experimental Manipulations and Prediction of Learning Outcomes. (A) Effects of experimental manipulations on Hidden Markov Model (HMM)-derived optimal K values. (B) Impact of experimental manipulations on p-values from inter-subject correlation analyses. (C) Association between K values and subsequent memory performance, quantified as Remember Score, within the *Commercial Movie Dataset*. (D) Association between K values and detailed subsequent memory performance, quantified as Detail Score, within the *Commercial Movie Dataset*. (E) Replication analyses examining the association between K values and memory performance, quantified as Remember Score, *in the STEM Course Dataset*. (F) Replication analyses examining the association between K values and memory performance, quantified as Test Score, in the *STEM Course Dataset*.

We then examined whether HMM-based event segmentation measures, specifically optimal K values and p-values, could predict learning outcomes in both the *Commercial Movie* and *STEM Course datasets*. For the *Commercial Movie Dataset*, memory performance for each participant was quantified using “remember” and “detail” scores from their speech-based free recall recordings. Across all participants, we found that the optimal K value was not associated with the number of events recalled (i.e., “remember” Score) (F=0.23, *p*=0.87; **Figure6C**), but was associated with the number of details recalled (“detail” Score) (F=7.36, *p*<0.001; **Figure6D**). The optimal K value was negatively correlated with the detail score (r=-0.42, *p*<0.001). No relationships were found between p values and memory performance (remember score: r=-0.07, *p*=0.45; detail score: r=0.04, *p*=0.63). For the *STEM Course dataset*, memory performance for each participant was quantified using “remember” and “test” scores. Across all participants, we found that the optimal K value was not associated with the Remember Score (F=1.02, *p*=0.39; **Figure 6E**) but was marginally associated with the test score: test scores decreased as the K value increased (r=-0.34, p=0.067; **Figure 6D**). We further found that as the p-value decreased, participants demonstrated higher test scores (r=-0.35, *p*=0.058), but not remember scores (r=-0.15, *p*=0.42).

Additionally, we assessed whether predictive effects were limited to HMM-driven event segmentation measures by examining correlations between *pupil size* and *moving speed* and memory performance. In the *Commercial Movie Dataset*, moving speed did not correlate with remember (r=-0.02, *p*=0.81) or detail score (r=-0.15, *p*=0.11). Pupil size did not correlate with remember score (r=-0.02, *p*=0.81), but negatively correlated with detail score (r=-0.21, *p*=0.032). In the *STEM course dataset*, neither the speed of movement (remember score: r=0.08, *p*=0.66; test score: r=0.20, *p*=0.28) nor pupil size (remember score: r=-0.19, *p*=0.31; test score: r=-0.22, *p*=0.24) showed significant correlations with learning outcomes. Consequently, we conclude that metrics derived from HMM event segmentation model, as opposed to simpler indices such as movement speed and pupil size, provide a stronger predictive value for learning outcomes as assessed by memory tests following video-based learning.

### 3.6 Results remain robust after correction for low-level confounds and gaze positions

To ensure that our findings reflected cognitive event segmentation rather than low-level sensory or motor artifacts, we performed a series of control analyses. We used a corrected pupil-size signal, from which potential confounds such as luminance and gaze position were regressed out, to re-evaluate our primary results. The main findings from the *Commercial Movie dataset* remained robust, whereas a few effects in the *STEM Course dataset* were no longer significant.

First, we re-examined pupil dilation and eye movement speed around event boundaries (Figure S3). In the *Commercial Movie dataset*, both pupil size (F(12, 1236) = 3.02, *p* < 0.001) and *eye movement speed* (F(12, 1236) = 87.808, *p* < 0.001) significantly increased, peaking at the boundary. A similar pattern was observed in the *STEM dataset* for both pupil size (F(12, 324) = 5.811, *p* < 0.001) and speed (F(12, 336) = 32.087, *p* < 0.001). Analysis of boundary strength in the *STEM dataset* revealed that pupil size was significantly modulated by high-strength (F(12, 24) = 1.562, *p* = 0.042) and medium-strength (F(12, 324) = 2.990, *p* < 0.001) boundaries, but not low-strength ones (F(12,324)=1.550, *p* = 0.105). In contrast, eye movement speed was sensitive to high-(F(12, 24) = 2.571, *p* < 0.001) and low-strength (F(12, 336) = 2.824, *p* = 0.001) boundaries, but not medium-strength ones (p = 0.168).

Next, we reassessed eye movement pattern similarity (Figure S4). For the *Commercial Movie dataset*, pattern similarity—incorporating both gaze and the corrected pupil signal— remained significantly higher between events than within events (F(2, 206) = 10.151, p < 0.001). This effect was not replicated in the STEM dataset, where no significant difference between conditions was found (F(2, 4) = 0.437, *p* = 0.782).

Similarly, we re-ran the ISC analyses using the corrected pupil data (Figure S5). In the movie dataset, ISCs remained significantly different across conditions (F(2, 206) = 44.66, *p* < 0.001), and boundary ISC was modulated by boundary strength (F(2, 84) = 17.26, *p* < 0.001). These effects were not observed in the STEM dataset (F(2, 56) = 2.20, p = 0.1208).

Lastly, we re-evaluated the HHM performance (Figure S6). For the *Commercial Movie dataset*, within-event similarity based on human annotations remained significantly higher than the null distribution (*p* < 0.001), whereas similarity based on HMM-generated boundaries did not (*p* = 0.507). Nevertheless, the HMM-generated boundaries aligned with human-annotated boundaries at an above-chance level (*p* = 0.009). For the *STEM dataset*, within-event similarity was significant for both human-annotated (*p* < 0.001) and HMM-generated boundaries (*p* < 0.001) compared to their respective null models. However, the alignment between HMM-generated and human-annotated boundaries did not reach significance (*p* = 0.078). A finer-grained analysis revealed this depended on boundary strength: the HMM identified high-strength boundaries (*p* = 0.04) but not medium-(*p* = 0.161) or low-strength boundaries (*p* = 0.654).

## Discussion

Using eye-tracking data from two distinct naturalistic viewing contexts—specifically, the viewing of a commercial movie and a STEM learning video—we demonstrated the efficacy of eye-tracking in measuring event segmentation during continuous naturalistic viewing. Our analysis began with basic metrics of *pupil size* and *movement speed*, and progressively incorporated advanced computational techniques, including *similarity analyses*, *intersubject correlation* (ISC), and *Hidden Markov Model* (HMM) analyses. Notably, these computational markers of event segmentation responded to event boundaries, were sensitive to experimental manipulations, and were able to predict learning outcomes effectively. Therefore, we concluded that eye-tracking offers sensitive indicators of event segmentation, with eye movement dynamics showing systematic relationships to event boundaries. Moreover, the integration of eye-tracking with neuroscientific techniques such as fMRI and EEG, alongside its applicability using common devices like laptops or smartphones, not only sheds new light on the fundamental theories of event segmentation but also holds substantial implications for industries like film production and online education.

Our initial investigation focused on the ability of basic eye-tracking metrics, specifically pupil size and movement speed, to track event boundaries during naturalistic viewing. We discovered that movement speed, in contrast to pupil size, serves as a more sensitive and reliable indicator for segregating events across movie and STEM learning video. Historically, pupil size has been a metric for assessing fluctuations in arousal related to ongoing cognitive and memory processes, serving as an insight into the neural underpinnings of cognition(Joshi & Gold, 2020; Sara, 2009). Relevant to the current study, there is growing evidence that arousal responses are attuned to the structure of temporally unfolding experiences, potentially indicating the event segmentation(Bianco et al., 2020; Clewett et al., 2020; Zhao et al., 2019). However, studies in this area have typically used auditory cues to demarcate event boundaries, which, although present, do not represent the primary method of segmenting information flow in the films and STEM materials in our study. Given the rich visual context of our naturalistic viewing conditions, we opted for movement speed as an alternative method for tracking event segmentation. This approach has been validated across two independent datasets and has demonstrated the modulatory effects of boundary strength on movement speed near event boundaries. In addition, we argue that movement speed is a more apt metric for measuring event segmentation during naturalistic viewing tasks, as it is less influenced by environmental factors such as lighting(Choi, 2017) or the participants’ internal states, including emotions and stress levels(Partala & Surakka, 2003).

Our pattern similarity analyses of eye movements (i.e., comparison between within and across event boundaries), revealed a notable reversal compared to a previous EEG study: pattern similarity was higher across event boundaries than within a single event. In contrast, Silva and colleagues, using EEG pattern similarity analyses, found that similarity within events exceeded that between events. They interpreted this finding as indicative of changes in neural representations upon detecting boundaries(Silva et al., 2019). Their finding was interpreted as indicating that neural representations of events change upon the detection of boundaries. The increased similarity around boundaries suggests its potential as an additional eye-tracking-based marker for event segmentation, although the underlying cognitive mechanisms remain unclear. The discrepancy between EEG and eye-tracking patterns warrants further investigation, possibly by applying the same analyses to fMRI data or by conducting simultaneous EEG and eye-tracking recordings during conditions of continuous video viewing.

Importantly, our ISC analyses of eye movements provided evidence that participants’ eye movements were more synchronized at event boundaries than during the event, suggests a strong link between collective information processing and event segmentation. It is a well-documented phenomenon that shared experiences can synchronize gaze positions and pupil sizes among individuals(Hasson, Landesman, et al., 2008; Madsen et al., 2021; Madsen & Parra, 2022). The shared neural and biological responses, captured by ISC and event segmentation, represent two major cognitive processes during naturalistic conditions, such as watching a movie or listening to music. However, few studies have linked these processes. One example could be: Clara and colleagues investigated variations in neural event boundary locations among individuals and demonstrated that those with more similar neural event boundaries in certain regions during movie-watching tended to have more similar movie appraisals(Sava-Segal et al., 2023). Our eye-tracking data provide direct evidence that eye movements were more synchronized at event boundaries, thus potentially serving as a major source of intersubject similarity. Considering the role of intersubject correlation and event segmentation in naturalistic perception and memory, we propose that the previously reported higher ISC values may predict successful memory formation(Gu et al., 2024; Hasson, Furman, et al., 2008; Madsen et al., 2021), at least partly, through enhanced event segmentation mechanisms. While one interpretation is that boundary perception causes eye movements’ synchrony, the relationship is likely more nuanced and potentially bidirectional. Increased gaze similarity might not just result from perceiving boundaries but could also contribute to their perception.

For instance, convergent fixation on salient transitional cues, leading to shared attention and increased inter-subject gaze synchrony, could heighten the collective likelihood of perceiving an event boundary. Furthermore, gaze synchrony might reflect shared predictions or anticipatory processes for upcoming transitions, sensitizing individuals to detect boundaries. Future research should aim to disentangle these intertwined influences on the dynamic interplay between shared gaze and event perception.

Inspired by the application HMM in neuroimaging(Baldassano et al., 2017), our study is the first investigation to whether determine if HMM can delineate event structure representation through eye-tracking data. Cross two distinct video stimuli (i.e., commercial movie and STEM educational video), our findings indicate a significant correspondence between event boundaries identified by eye-tracking & HMM and those annotated by humans. This alignment echoes the alignment observed in studies employing fMRI(Baldassano et al., 2017; Geerligs et al., 2021, 2022; Yates et al., 2022) and EEG(Silva et al., 2019; Sols et al., 2017), and the alignment led to significant higher similarity values in the eye-tracking data. Given the affordability and accessibility of eye-tracking technology, this confirmation opens new avenues for future research that might integrate neural-based event segmentation studies with eye-tracking methodologies. Importantly, our analysis revealed considerable variability in the concordance between HMM model predictions and human annotations, as well as the influence of boundary strength (i.e., event boundaries with higher strength are more easily identified by the HMM). The variation in HMM predictions should not be dismissed as mere noise but rather considered as a potential objective indicator for event segmentation, extending beyond traditional neural markers such as hippocampal responses(Ben-Yakov et al., 2013; Ben-Yakov & Henson, 2018; Cohn-Sheehy et al., 2021; Reagh et al., 2020). While our primary analyses aimed at conceptually meaningful event segmentation with number of events closely reflecting human annotation, we also explored finer-grained segmentations using HMMs with substantially larger event counts (100+). These high-resolution segmentations primarily reflected rapid attentional shifts rather than meaningful conceptual boundaries, showing weaker relationships with learning outcomes. Thus, while finer-grained approaches offer detailed attentional insights, our current conceptual-level approach aligns better with existing cognitive and neural frameworks suggesting segmentation at broader, cognitively meaningful temporal scales.

Employing data from experimental manipulations preceding video viewing and subsequent memory assessments, we conducted a comprehensive examination of how pupil diameter, movement speed, k value from HMM, and the P-value of similarity analyses vary as a function of experimental conditions and their predictive capability regarding learning outcomes. Previous studies have indicated that factors such as development(Yates et al., 2022), age(Reagh et al., 2020), anticipation(Lee et al., 2021), and schema(Masís-Obando et al., 2022) can influence fMRI markers of event segmentation. Our results suggest that compared to pupil size and moving speed, metrics derived from HMM display enhanced sensitivity to various psychological states, potentially augmenting or impairing event segmentation. This insight is pivotal for investigating event segmentation across different demographics and experimental conditions in future studies. Eye-tracking-based ISC has been recognized for its ability to predict learning outcomes(Gu et al., 2024; Madsen et al., 2021; Madsen & Parra, 2022). Here, we augment the predictive scope of eye-tracking metrics by demonstrating that metrics associated with HMM event segmentation model can also foresee subsequent memory performance. Unlike ISC analyses, whose underlying cognitive theories remain ambiguous, event segmentation has long been hypothesized to play a critical role in memory formation(Kurby & Zacks, 2008; Zacks, 2020). The eye-tracking-based event segmentation index offers a more interpretable metric for delineating underlying mechanisms and identifying targets for specific cognitive and/or neural modulation-based learning strategies. Why does the event segmentation-based index, as opposed to merely pupil size and movement speed, predict learning outcomes? We examined the sensitivity of our HMM-based eye-tracking approach to different viewing conditions by including videos varying substantially in their narrative structure and pacing. While prior studies have shown minimal effects of strong top-down comprehension manipulations or reversed playback on event segmentation and eye movements (Hard et al., 2006; Huff et al., 2017; Hutson et al., 2017), we propose that the rapidly-paced, fragmented nature of short-form content may uniquely influence event segmentation processes. Specifically, short-form video viewing might induce distinct patterns of eye movements, characterized by increased saccade frequency and altered fixation durations, which our HMM approach is particularly sensitive in detecting. Supporting this interpretation, the optimal number of events identified by the HMM (k-value) differed across video conditions and correlated meaningfully with memory performance: higher fragmentation (higher k-value) was associated with poorer memory recall. Thus, our findings extend the current understanding of event segmentation, demonstrating the practical utility and sensitivity of eye-tracking-based HMM analyses for capturing subtle cognitive effects induced by diverse real-world viewing conditions.

A critical consideration for pupillometry in naturalistic settings is the influence of non-cognitive factors, as pupil size is highly sensitive to low-level visual features (e.g., luminance, contrast) and gaze position. To ensure our findings were not driven by such stimulus-related confounds, we implemented a correction procedure based on the methods of Antony et al. (2021) to regress out these influences. Our core findings remained highly robust following this correction. Specifically, the Hidden Markov Model (HMM) trained on the corrected pupil data continued to reliably identify event boundaries that aligned with human annotations and retained its sensitivity to experimental manipulations. This result confirms that robust cognitive signals of event segmentation can be extracted from pupillary data. However, the application of such corrections involves a trade-off. While computationally intensive, this step provides methodological rigor by isolating cognitive signals. We advocate for future research to carefully consider this balance between rigor and practical feasibility, especially in contexts involving real-time analysis or resource-constrained settings.

Our results replicate and extend prior findings regarding eye movement behaviors at event boundaries. Like Eisenberg’s study(Eisenberg & Zacks, 2016), we observed dynamic alterations in eye movements around event boundaries, including increased movement speed and pronounced pupil dilation, reinforcing the idea that eye movement behaviors reflect cognitive transitions during segmentation. Moreover, consistent with recent findings from two previous studies (E. E. Davis et al., 2021; Smith et al., 2024), we observed heightened inter-subject correlations at event boundaries, underscoring the synchronized nature of visual attention during significant event transitions. Notably, our application of Hidden Markov Models (HMM) adds a novel methodological dimension, demonstrating the predictive capability of computationally derived event boundaries for learning outcomes. Thus, our study integrates and expands upon previous literature by providing a comprehensive computational framework that advances our understanding of event segmentation as measured through naturalistic eye-tracking methods. Our investigation is the initial endeavor to measure event segmentation through eye-tracking data, with certain limitations noted. Firstly, event segmentation’s occurrence across diverse video stimuli is acknowledged. Our analysis of two distinct eye-tracking datasets—captured while participants viewed either a commercial movie or a STEM course video—aims to broaden our study’s applicability and act as a conceptual replication. They represent typical daily activities but possess distinct editing features (e.g., cuts, scene changes, switch of slides) that may facilitate event segmentation detection through eye-tracking(Magliano et al., 2020; Magliano & Zacks, 2011). Conversely, everyday activity videos, like personal vlogs (e.g., purchasing coffee at Starbucks), may lack these editing traits, possibly limiting eye-tracking’s efficacy in event segmentation analysis. Secondly, our data were gathered advanced eye-tracking technology within controlled laboratory settings to ensure data quality. Nevertheless, the application of event segmentation identification in real-world settings might rely on data from cost-effective cameras and less regulated environments, posing challenges that future research must address. A notable study used ISC to analyze eye movements captured by standard web cameras in classroom and home setting, and successfully predicted tests scores during online education(Madsen et al., 2021). This study may suggest that eye movement metrics might be robust against variations in measurement tools and environmental disturbances compared to neuroimaging benchmarks. Thirdly, our participant pool consisted of healthy young adults from a university setting, generally exhibiting good attention and comprehension of the videos. While event segmentation has been effectively studied in other demographics, including older adults(Reagh et al., 2020) and infants(Yates et al., 2022), eye tracking data from a broader demographic spectrum may exhibit higher blink rates and tracking loss, adversely affecting measurement accuracy(Dalrymple et al., 2018; R. Davis, 2021). We argue that research extending to diverse groups or psychiatric patients should consider extended recording times or increased sample sizes to effectively analyze event segmentation processes through eye-tracking. Another potential limitation of our eye-tracking methodology is the use of subtitles in movie stimuli, which likely influenced participants’ gaze patterns by drawing attention towards the lower part of the screen. Although our analytical methods focus primarily on dynamic eye movements and pupil size changes indicative of cognitive event processing rather than absolute gaze fixation points, future research could explicitly quantify subtitle-related gaze distribution effects. Replication studies using materials without subtitles or systematically varied subtitle conditions could further validate the robustness and generalizability of eye-tracking methods in measuring event segmentation.

In conclusion, eye-tracking offers an alternative methodology for researchers investigating the event segmentation process during naturalistic video viewing. This approach can be employed concurrently with other neuroscientific techniques, such as EEG and fMRI, to explore eye-brain interactions during event segmentation. Furthermore, it can be used in online experiments using laptop or smartphone cameras to enable large-scale studies of event segmentation in real-world settings.

## Declarations

• **Funding** (information that explains whether and by whom the research was supported) W.L was supported by the National Natural Science Foundation of China (grant No. 32300879), Humanities and Social Sciences Fund, Ministry of Education (grant No. 22YJCZH109), Natural Science Foundation of Hubei (grant No. 2022CFB793). X.H was supported by the Knowledge Innovation Program of Wuhan-Shuguang Project (2023020201020384).

• **Conflicts of interest/Competing interests** (include appropriate disclosures)

The authors declare that they have no known competing financial interests or personal relationships that could have appeared to influence the work reported in this paper.

• **Ethics approval** (include appropriate approvals or waivers)

Experimental procedures (both Study1 and Study2) were approved by the University Institutional Review Board for Human Subjects.

• **Consent to participate** (include appropriate statements)

All participants (both Study1 and Study2) provided written informed consent before the start of experimental procedure.

• **Consent for publication** (include appropriate statements)

All authors read the paper and povided consent to submit it to Behavioral Research Methods

• **Availability of data and materials** (data transparency)

Processed behavioral and eye-tracking data, as well as custom-made video stimuli, are available at OSF (https://osf.io/n7k45/?view_only=673d89b23720410799b5daec1f5d511e)

• **Code availability** (software application or custom code)

Scripts for performing all eye-tracking analyses are located in the same repository as the data and stimulia at OSF (https://osf.io/n7k45/?view_only=673d89b23720410799b5daec1f5d511e)

• **Authors’ contributions** (optional: please review the submission guidelines from the journal whether statements are mandatory)

W.L. and X.H conceived the Study; J.S.L and Z.Y.C collected the data; W.L. and J.S.L analyzed the data; W.L., X.H, and J.S.L prepared the first draft. W.L. and X.H reviewed and edited the manuscript, provided supervision, and obtained funding.

