## Supplemental files for "Boundaries in the eyes: measure event segmentation during naturalistic video watching using eye tracking"

### for

**This PDF file includes:**

Figure S1-S6

**Correspondence:**

Dr. Xin Hao

School of Psychology, Central China Normal University

Address: Nanhu Complex Building, No 152 Luoyu Road, Wuhan. Postal Code: 430079

Dr. Wei Liu

School of Psychology, Central China Normal University

Address: Nanhu Complex Building, No 152 Luoyu Road, Wuhan. Postal Code: 430079

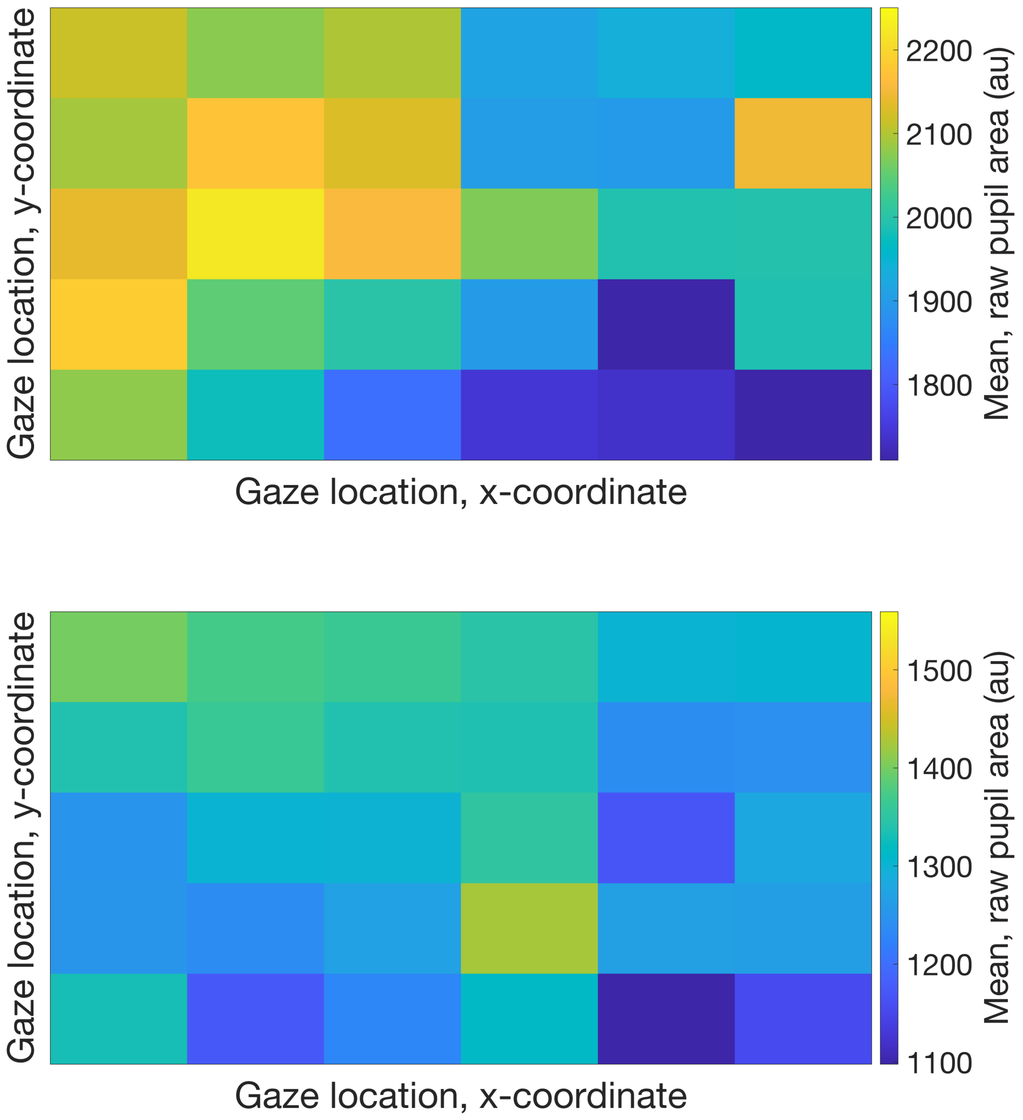


**FigureS1 Pupil size as the function of Gaze location in the movie dataset (up) and in the STEM dataset (down)**

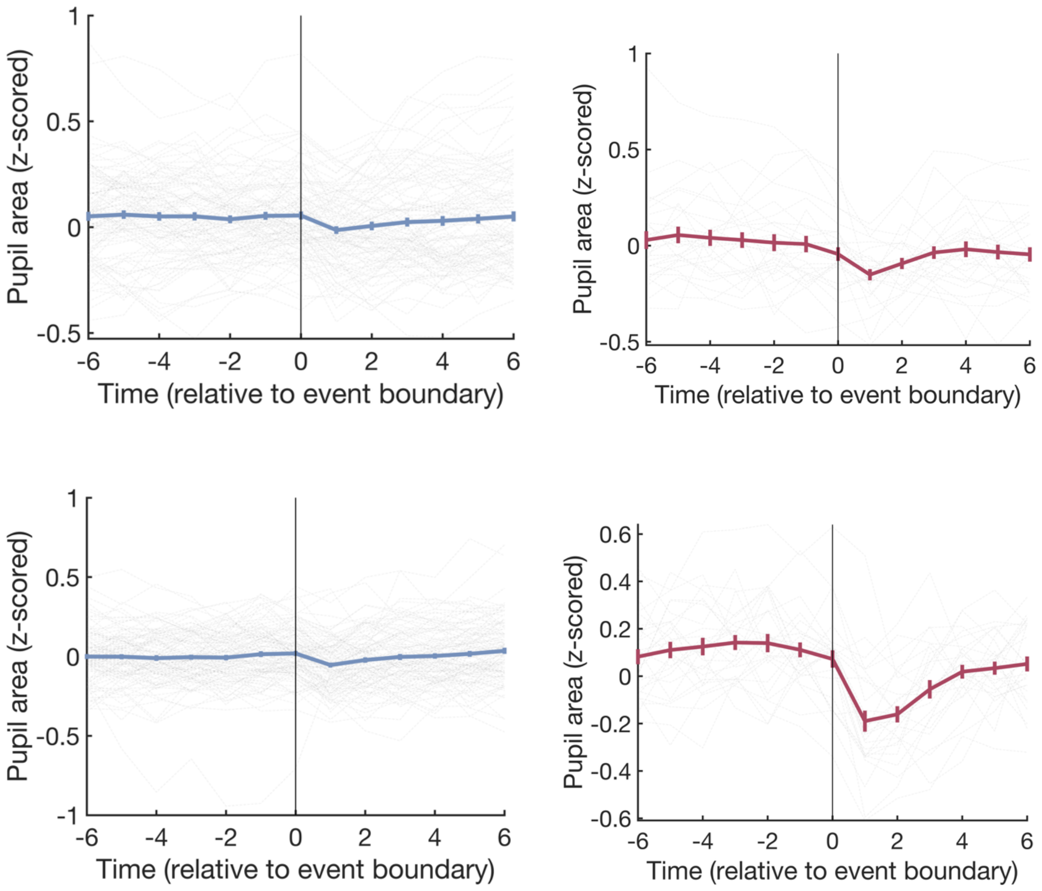


**FigureS2 Uncorrected Pupil size as the function of event boundary in the movie dataset (up-left) and in the STEM dataset (up-right). Corrected Pupil size as the function of event boundary in the movie dataset (down-left) and in the STEM dataset (down-right)**

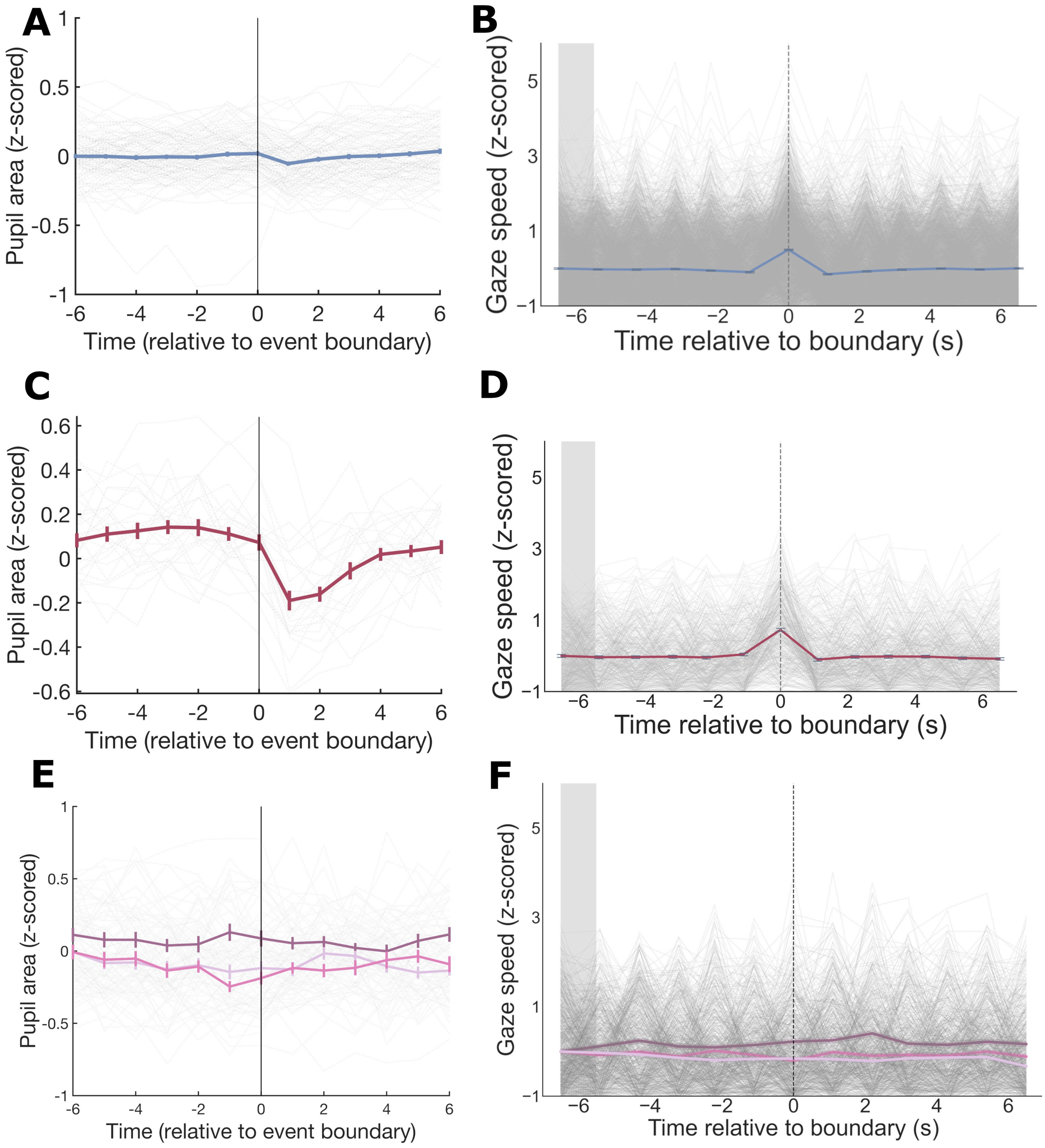


**FigureS3 (corresponding to main Figure2, analyses were performed with corrected pupil area). Pupil Size and Eye Movement Speed in Response to Event Boundaries and Their Modulation by Boundary Strength.** (A) Pupil size dynamics around event boundaries in the Commercial Movie Dataset. (B) Eye movement speed dynamics around event boundaries in the Commercial Movie Dataset. (C) Pupil size changes around event boundaries in the STEM Course Dataset. (D) Eye movement speed dynamics around event boundaries in the STEM Course Dataset. (E) Impact of event boundary strength on pupil size adjustments in the Commercial Movie Dataset. (F) Influence of event boundary strength on eye movement speed patterns in the Commercial Movie Dataset.

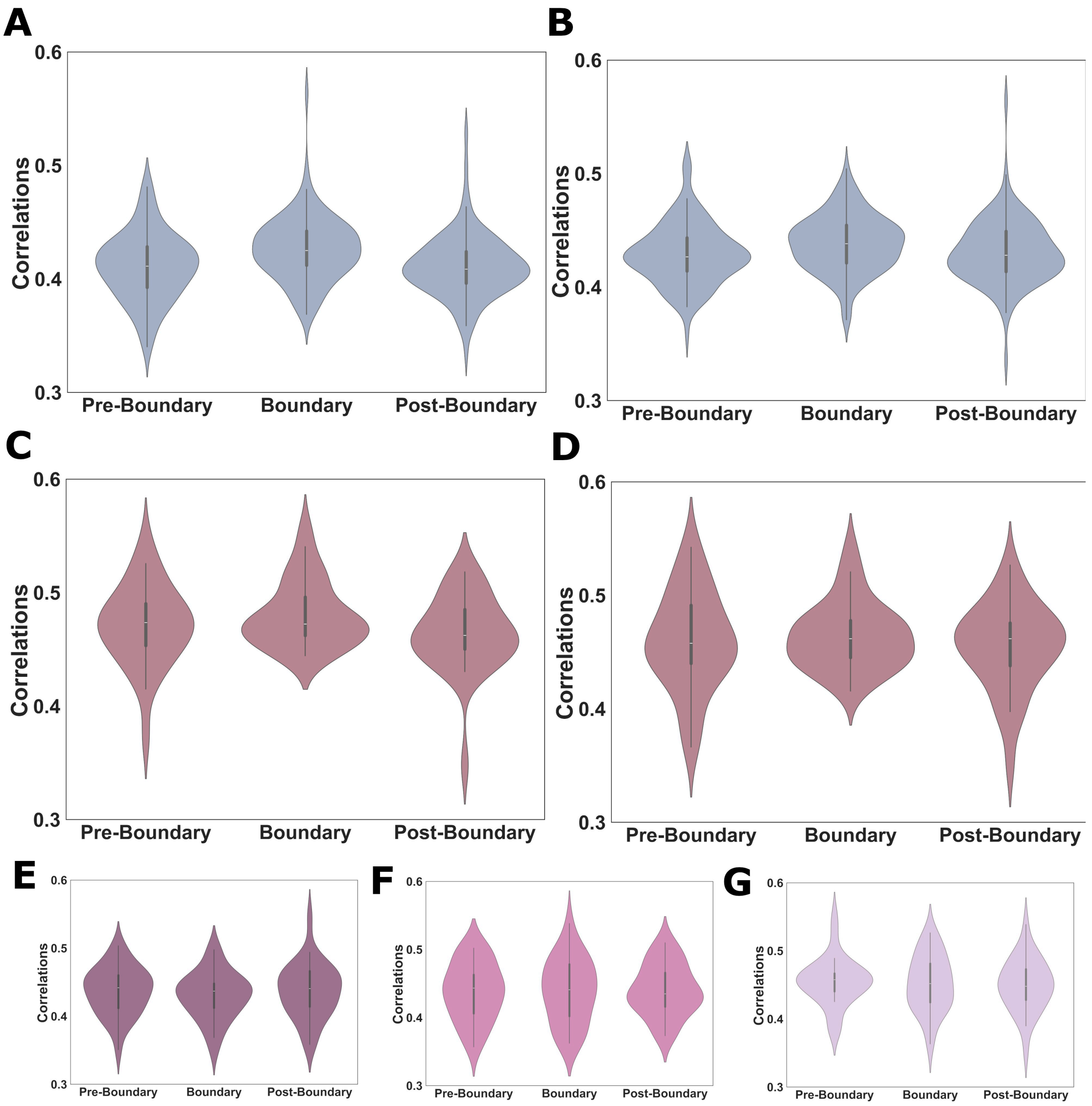


**FigureS4 (corresponding to main Figure3, analyses were performed with corrected pupil area). Eye movement pattern similarity within and across event boundaries during continuous video watching.** (A) We analyzed three similarity measures derived from pupil size—within events before the boundary, across events separated by boundaries, and within events after the boundary—using the Commercial Movie Dataset. (B) Higher similarity measure, incorporating both pupil size and vertical and horizontal gaze, was calculated in the across-event condition, compared to two within-event conditions, using the Commercial Movie Dataset. (C) Participant's three similarity measures derived from pupil size for three event conditions, analyzed with the STEM Course Dataset. (D) Participants' similarity measures, calculated from both pupil size and vertical and horizontal gaze, were evaluated for three event conditions with the STEM Course Dataset. (E–G) Combined similarity measures (pupil size and gaze position) stratified by event-boundary strength—high (E), medium (F), and low (G)—in the STEM Course Dataset.

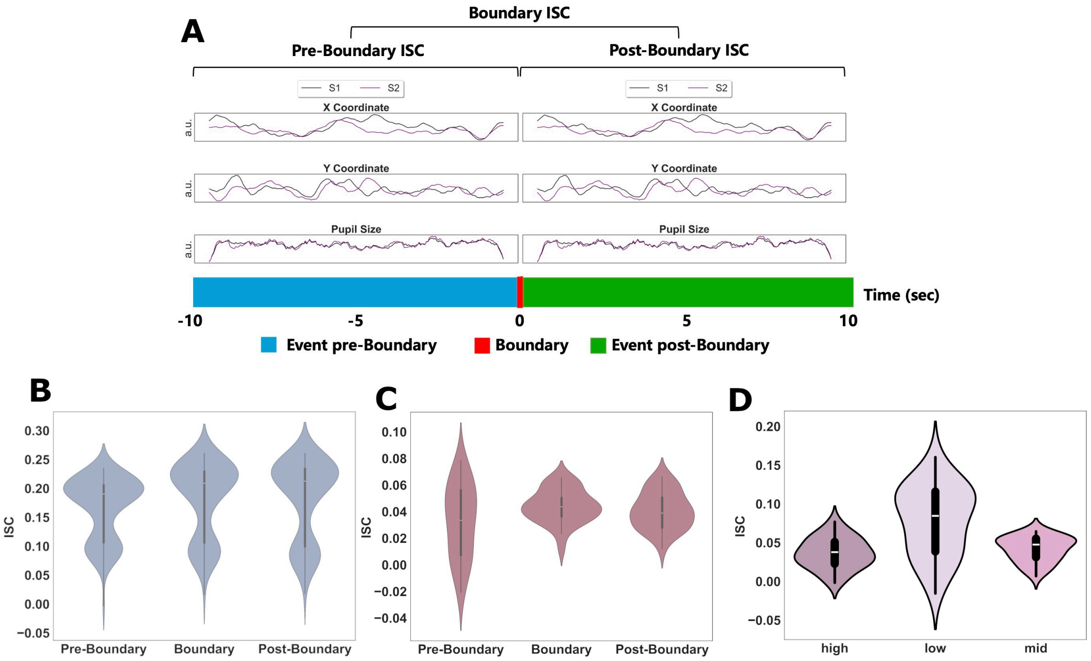


**FigureS5 (corresponding to main Figure4, analyses were performed with corrected pupil area). Synchronized Eye Movements in Response to Event Boundaries.** (A) Synchronized eye movements—specifically, the positions (X and Y coordinates) and pupil sizes—of two participants (S1 and S2) while continuously watching two distinct events separated by an event boundary. Intersubject correlation (ISC) is quantified as the mean correlation between both vertical and horizontal gaze positions and pupil sizes across participants. We analyze three ISC metrics: pre-boundary ISC, calculated during the final 10 seconds before the event boundary; boundary ISC, derived from 5 seconds of eye-tracking data from each event; and post-boundary ISC, measured during the initial 10 seconds following the event boundary. (B) Comparison of three kinds of ISC within the Commercial Movie Dataset. (C) Comparison of three kinds of ISC within the STEM Course Dataset. (D) Comparison of boundary ISC associated with event boundaries characterized by high, medium, and low boundary strengths.

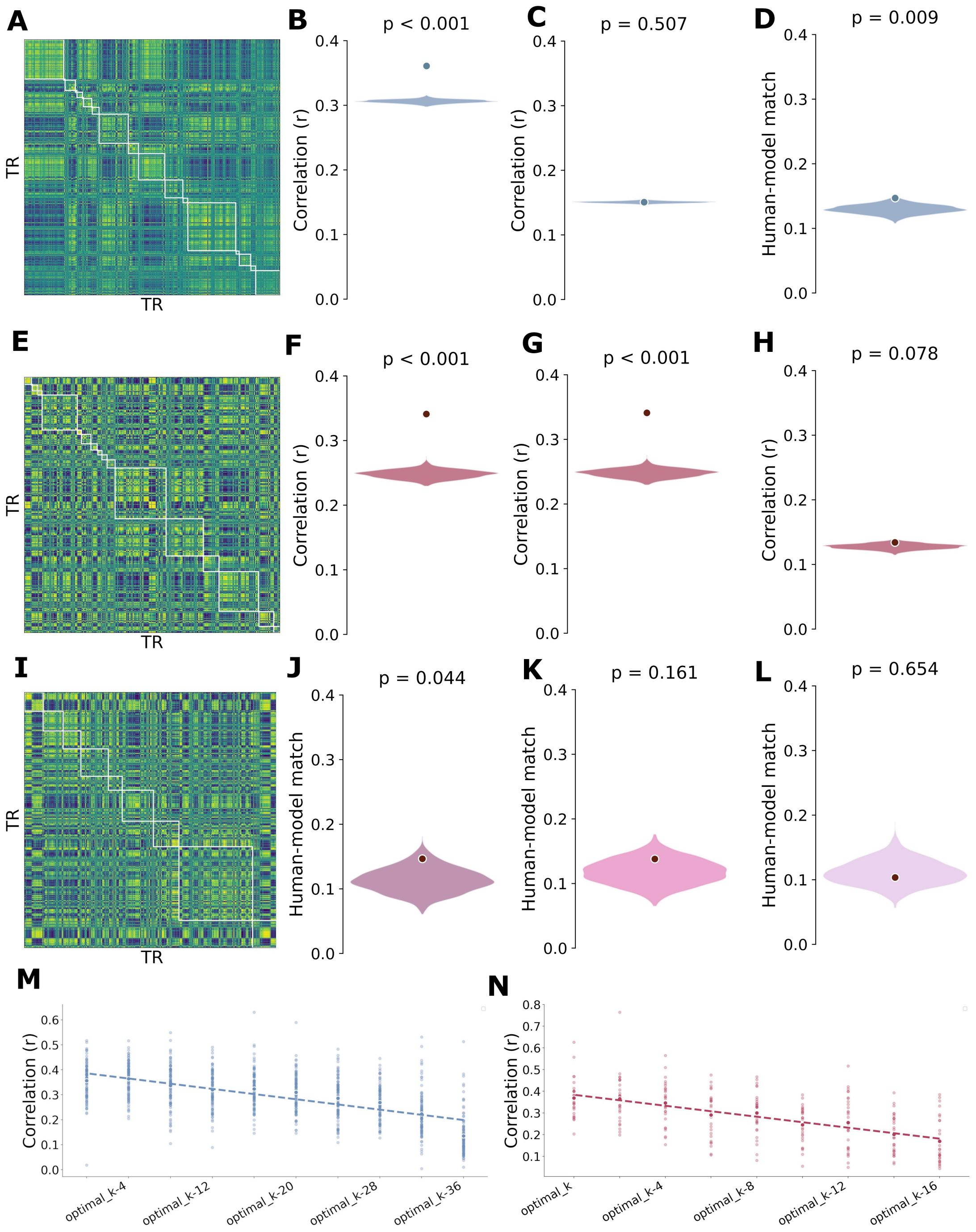


**FigureS6 (corresponding to main Figure5, analyses were performed with corrected pupil area). Hidden Markov Model (HMM)-based Event Segmentation Model for Eye-Tracking Data.** Analyses on the Commercial Movie dataset (A-D, M): (A) Example alignment of HMM and human-annotated event boundaries. (B-C) Within-event gaze similarity was significantly higher than the null distribution (histograms) for both (B) human-annotated and (C) HMM-generated boundaries. The light blue circle indicates the observed participant average. (D) HMM-human boundary matching was significantly above chance. Corresponding analyses on the STEM Course dataset (E-L, N): (E) Example boundary alignment. (F-G) Within-event similarity for (F)

human and (G) HMM boundaries. (H) HMM-human boundary matching. Boundary strength and model parameter analyses: (I) Example alignment for a high-strength boundary in the STEM dataset. (J-L) Boundary matching was modulated by strength, with the HMM effectively identifying high-strength but not medium- or low-strength boundaries. (M-N) The relationship between the HMM's optimal event count (K) and gaze similarity is shown for the (M) Commercial Movie and (N) STEM Course datasets.
